# Clones on the run - the genomics of a recently expanded facultative asexual species

**DOI:** 10.1101/2022.05.11.491277

**Authors:** Ricardo T. Pereyra, Marina Rafajlović, Pierre De Wit, Matthew Pinder, Alexandra Kinnby, Mats Töpel, Kerstin Johannesson

## Abstract

Why, in facultative asexual species, marginal populations are often richer in clones than are core populations, remains unclear. Cloning freezes genotypes but hampers recombination and local adaptation. During expansion, clones are favoured over non-selfing sexuals by uniparental reproduction. To better understand the dynamics of clones and sexual lineage, we used genome-wide sequencing to analyse a recently expanded seaweed. We found large clones and sexual populations mixed close to range margins. Clones had evolved repeatedly from sexual populations but were unexpectedly low in genetic variation. Modelling suggested clones form from sexual populations after repeated bottlenecks at the expansion front. A clonal wave of depauperate genotypes thereafter spread ahead of the sexual population. As we observed, these early formed clones may survive side-by-side sexual individuals, which suggests they lost their sexual capacity. Our study illustrates how range expansion can result in complex and dynamic patterns of genetic variation in facultative asexual species.

**Teaser:** We use genome data and modelling to find out why large clones are only found at range margins in a recently expanded seaweed

## Introduction

Range expansion following retreat of the northern ice-cap (<15,000 years ago) shaped the demographic history of species in the northern hemisphere that still impacts their genetic structure (1, 2). Lessons from postglacial expansions can help us understand effects of species range modifications under current climate change including introductions of alien species into new areas. Key questions include the nature of genetic processes at the front of the expansion (3-5), the role of recombination versus asexual reproduction at range margins (6-9), and patterns of dispersal including the role of long-range dispersal (10-12).

Expansion processes have been described in various systems and many notoriously feature clonality as a determinant for the colonisation in otherwise mostly sexual species (13-15). Clonality is a generic trait promoting rapid expansion in many eukaryote taxa (16-21), as well as in organoids (22), cancer cells (23), bacteria (24), and parasites (25). Many species of plants, seaweeds, and invertebrates can reproduce both sexually and asexually, and these facultative asexual species tend to show a branching phylogenetic pattern - known as a “twiggy” distribution (26), where the expanding clonal lineages have relatively short branches derived from and spread among sexual counterparts. Such an array of lineages suggests a limited evolutionary lifespan of clones, while clones can still be dominant during thousands of years. The presence of clones in a facultative asexual species can have various effects on the genetic structure of the species (27). For example, while drift rapidly reduces genetic variation in small and fragmented populations at the expansion front (4), in clones single-locus heterozygosity is more readily accumulated, by new mutations, and protected. Accumulation of new somatic mutations also gives rise to segregation of new clones, followed by a potential for drift and selection among clones of a species (21). Patterns of expansion in facultative clonal species have been described mostly in short-generation model systems, or in ancient clonal species. In the latter, the expansions occurred a very long time ago and details of the initial colonisation phase are difficult to disentangle. Understanding the early phases of colonisation and expansion into new territory in long-lived non-model organisms with the capacity for both sexual and asexual recruitment remains an urgent task. This includes tracing the sources of genetic variation of the expanding population, and how this variation evolves and becomes spatially structured. However, few such expansions have been characterised and the details remain largely elusive.

*Fucus vesiculosus* is a foundation species of north Atlantic shallow rocky areas, and a main component of “marine forests” of coastal areas. In large parts of the Baltic Sea (a marginal brackish-water basin connected to the NE Atlantic through the Danish Straits) it is the only perennial macroalgae, and as such, a key habitat for marine invertebrates and fish (28). The species is dioecious (separate male and female individuals), with eggs and sperm formed in receptacles at the frond tips and released during spring and summer. After release, gametes are short-lived and soon after fertilisation the egg becomes negatively buoyant, drops to the bottom, settles, and develops into a new diploid individual (29). Consequently, gametes and zygotes have a very restricted dispersal, but occasionally an individual can be wrenched loose during a storm and drift to a new area (30). Approximately 8000 years ago, following the retraction of the glacial ice sheet, a former large freshwater lake opened into the Atlantic and formed the Baltic Sea (31). Earlier studies suggest that *F. vesiculosus* expanded from the NE Atlantic into the Baltic Sea soon after the opening (32, 33). Despite being earlier considered an obligate sexual species it forms large clones in the Baltic Sea, and asexual reproduction is by shedding of adventitious branches that reattach to the rock surface (34). Individuals that recruit asexually are copies of sexually recruited individuals and form reproductive tissue and gametes (35). Asexual reproduction has been suggested unique to Baltic *Fucus* and a consequence of the low salinity of this postglacial basin (36-38).

We here examined the evolutionary history of *F. vesiculosus* throughout its establishment inside the Baltic Sea and outside in neighbouring parts of its NE Atlantic distribution. The expansion of *F. vesiculosus* from the Atlantic into the Baltic Sea also includes a recently (4000 y ago) diverged sister species, *F. radicans* endemic to the Baltic Sea (32, 39). The systematic relationship between the two taxa remains unclear (40) but as these taxa represent a monophyletic lineage expanding into the Baltic Sea (32) we here treat both as *F. vesiculosus*. We asked why asexual reproduction and clonal lineages have established during colonisation of this postglacial sea and how it affects the current genetic landscape. In an earlier study (11), using a mathematical model describing the evolution of clonal lineages, we made two predictions: First, uniparental advantage during colonisation of an empty area will favour asexual reproduction during range expansion leading to a clonal wave that spread ahead of a sexual wave. This will happen despite cloning being subordinate to sexual reproduction in core populations. Second, occasional long-distance dispersal will enhance encounters of opposite sex that result in new clones and multiple local clonal waves, and add genetic complexity to the system. In the present study we generated extensive genomic data in *F. vesiculosus* throughout the Baltic and adjacent NE Atlantic areas to evaluate these predictions. In addition, we extended the previous model and included descriptors of genetic variation and inbreeding to better understand the processes involved in the formation of the clonal waves, and the spatial and temporal dynamics of genetic variation.

## Results

We obtained a detailed description of the spatial pattern of genetic variation in *F. vesiculosus* by genome-wide genotyping of 897 individuals from 47 sites in the Baltic Sea and nearby areas of the Atlantic Ocean (Suppl. Table S1) using 2b-RAD and >15,000 SNPs (41). We applied three different descriptors of population genetic variation: *i*) Expected heterozygosity (*H*_e_) indicating overall genetic variation of a population, *ii*) observed heterozygosity (*H*_o_) that increases from the accumulation of new mutations in clonal lineages, and *iii*) the inbreeding index (*F*_IS_) as a main descriptor of clonality. In clonal lineages *F*_IS_ will decrease below zero and be close to -1 at complete clonality (42), but even low rates of recombination (in facultative asexual populations) and mutation tend to increase *F*_IS_ (43). Consequently, *F*_IS_ can be used as a sensitive indicator of low rates of recombination in a population that is mostly asexual (44). A low rate of recombination is potentially a very important feature of a facultative asexual population, as it will introduce new genotypes and promote divergence through formation of new clonal lineages. *F*_IS_ has a close relationship to observed and expected heterozygosity (*F*_IS_=1-*H*_o_/*H*_e_). In a large well-mixed sexual population, the expectation is that *H*_e_ and *H*_o_ are approximately equal and *F*_IS_ is close to zero. In populations that reproduce both sexually and asexually there is no similarly strong prediction (43). Not least will the realised overall proportion of local asexual reproduction vary both spatially and temporally in a facultative asexual species (11).

### The genomic landscape of *Fucus vesiculosus* in a marginal environment

Overall, we found a strong divergence among the 47 *Fucus vesiculosus* sites (global *F*_ST_= 0.073) and a spatial structure in which individuals almost exclusively clustered genetically according to their sampled site (Fig. 1A). This is consistent with a predominantly short-range dispersal. At a higher hierarchical level, the NJ tree revealed that all individuals sampled inside the Baltic Sea belonged to a separate Baltic *Fucus* cluster diverged from all individuals sampled in the Atlantic. Individuals from the “Transition zone”, at the entrance of the Baltic Sea, where the salinity shifts from <10‰ to >25‰ over 250 km (45), formed a basal group in the large Baltic Sea cluster. This overall pattern corroborates a postglacial colonisation of the Baltic Sea from the Atlantic through the Transition zone after this brackish water basin opened to the Atlantic 8000 years ago.

**Fig. 1.**
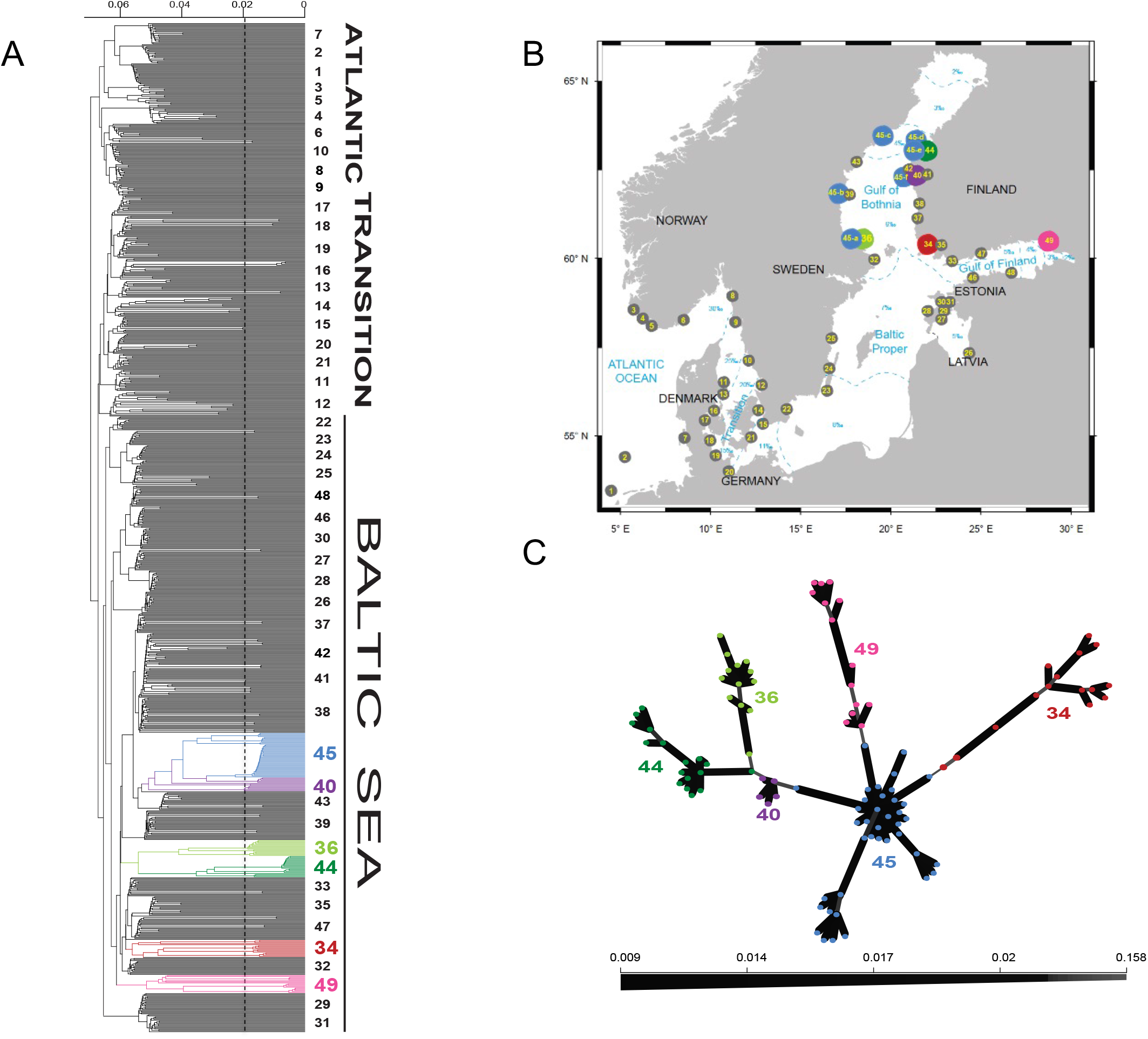
Study area and genetic relationships among *Fucus vesiculosus* sampled in 47 different sites. Site 1-Roscoff in France and site 2-Bangor in UK are outside the map. **A)** Neighbour-Joining Tree based on 30,581 2b-RAD loci showing the genetic relationships among samples. In most cases each “lineage” contained individuals of *F. vesiculosus* from only one site, except for lineage 45 that were present in 5 sites (denoted a-e). Separate clonal clusters (see text) are illustrated in different colours. Vertical dashed line indicate cut-off for identification of clones, see text. **B)** Map showing the geographic distribution of the sites and the salinity (‰) gradient from the Atlantic and into the Baltic Sea. **C)** Minimum Spanning Network showing inter- and intraclonal reticulations among clonal lineages based on genetic dissimilarity distances. Thickness of connecting internodal lines depict the magnitude of genetic distance between individual nodes as indicated in the lower bar

Most individuals in the NJ tree clustered by long branches (black in Fig. 1A), while individuals from eight sites clustered mainly by short branches (coloured branches in Fig. 1A and sites marked with large dots in Fig. 1B). Short branches potentially indicative of somatic mutations suggested the presence of clones (46-48), but the occurrence of intermediate length branches complicated the discrimination of clonal and sexual genotype groups. For this purpose, we used a genetic distance-based method with three different clustering algorithms (49 and see Methods) to infer the genetic distance threshold separating asexually recruited genotypes. While there was discrepancy in one clustering algorithm, the other two overlapped at a genetic distance of 0.02 (vertical line in Fig. 1A). Branches shorter than this distance had only minor genetic differences (presumably reflecting somatic mutations) and were considered as members of the same clone (multi locus genotype, MLG). Between three and eight clones grouped together into six different clusters (illustrated by separate colours in Fig. 1A). We hereafter refer to each such cluster as a “clonal cluster”.

The mix of clonal clusters and sexual populations in the NJ tree showed that the clonal lineages had evolved repeatedly from sexual lineages during the expansion into the Baltic Sea. We used a Minimum Spanning Network (MSN) to look further into the possibility of reticulation among clonal clusters instead of repeated independent evolution of these clusters. The MSN analysis, however, showed no reticulation among the major clusters, yet it corroborated close ancestries between cluster pairs, in particular clusters 40 and 45 (Fig. 1C). Interestingly, cluster 40, exclusively found in a Finnish site (Sälskär, Suppl. Table 1) appears basal to cluster 45 which is spread over several, mostly Swedish, sites (Fig. 1B).

**Table 1.** Model parameters. Notations used (first column), brief explanation (second column), and values explored (third column).

| Notation | Explanation | Values |
| --- | --- | --- |
| $K$ | Number of demes | 500 |
| $K_0$ | Number of source (core) demes | 50 |
| $N$ | Maximum number of individuals in each deme | 100 |
| $S$ | Per-male maximum number of sexual encounters | 8 |
| $C$ | Per-individual number of clonal offspring | 1 |
| $m_T$ | Per-individual total migration rate | 0.0001, 0.006, 0.05 |
| $p_L$ | Probability of long-range dispersal per migrant | 0, 0.001, 0.007 |
| $s_{Loss}$ | Per-individual rate of loss of sex | 0, 0.0005, 0.005 |
| $d$ | Per-individual death rate | 0.1 |
| $L$ | Number of loci | 100 |
| $\mu$ | Per-individual per-allele mutation rate | 0.000025 |
| $H_e^{(i)}$ | Starting per-locus $H_e$ in the source demes<br>$i = 1, 2, \dots, K_0$ | 0.2 |

While large clones and large clonal clusters were only found in the marginal areas of the Baltic Sea, pairs of clonemates were occasionally present throughout the study area (Fig. 1A). Such pairs were most frequent in the southern Baltic Sea and in the Transition zone, but also found in one Atlantic site (site 6), suggesting that asexual recruitment of individuals in *F. vesiculosus* is also present outside the Baltic Sea. Five of the clonal clusters had local distributions, but cluster 45, the most distal in the NJ tree, was widespread and found at six sites (45a-45f in Fig. 1B) over > 600 km of coastline. This wide distribution not only applied to the whole clonal cluster, but also to the most frequent clone inside this cluster (Suppl. Fig. S1).

### Spatial patterns of *H*_e_, *H*_o_, and *F*_IS_

To illustrate changes along the colonisation route in relative genetic variation (*H*_e_), heterozygosity (*H*_o_), and the inbreeding index (*F*_IS_) we plotted these variables along the NE Atlantic to Baltic Sea colonisation route (Fig. 2; the x-axis illustrating the order rather than the geographic positions along the route). Disregarding the clonal clusters (green dots in Fig. 2) both *H*_e_ and *H*_o_ tended to weakly decrease into the Baltic sites, but the differences between Atlantic and Baltic sites were minor, while the Transition zone peaked in both *H*_e_ and *H*_o_ relative to the Atlantic and Baltic sites (Fig. 2, Suppl. Table 2).

**Fig. 2.**
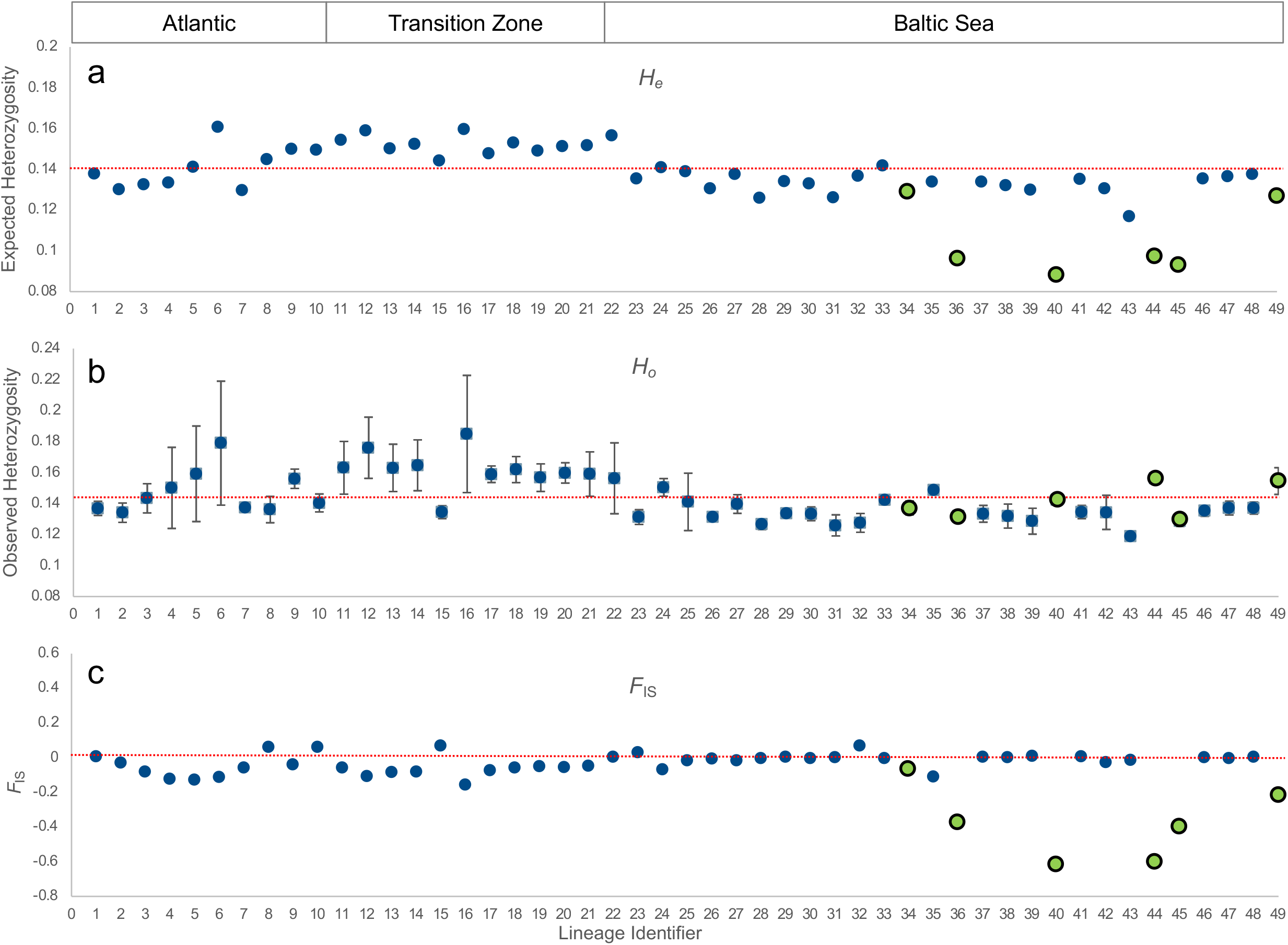
Variation in *H*_e_ (A), *H*_o_ (B) and *F*_IS_ (C) among sexual populations and clonal clusters. Please note that sample 45 were present in five sites. Sites are arranged along the x-axis (not to scale) starting from SW Atlantic and ending in the innermost parts of the Baltic Sea and site numbers are the same as in Fig. 1B. If the same site include one clonal cluster and one sexual population, or two clonal clusters, these have different numbers (see Suppl. Table 1). Green points represent clonal clusters, red dashed lines indicate overall averages of *H*_e_ and *H*_o_ (clonal clusters not included) and 0 of *F*_IS_, respectively. Error bars indicated for *H*_o_ illustrate SD. among the 30,581 2bRAD loci in each population (where clonal clusters are treated separately when sampled from the same sites as sexual populations).

As clonal lineages, according to evolutionary theory, are resistant to drift and prone to accumulation of new mutations, our prediction was that clones would contain slightly higher genetic variation and much higher observed heterozygosity (due to lack of segregation of homozygotes) than sexual populations. However, four of the clonal clusters had substantially lower *H*_e_ than sexual populations (Fig. 2A) and *H*_o_ values similar to nearby sexual populations (Fig. 2B). Finally, five out of six clonal clusters had more negative *F*_IS_ values than any of the sexual populations (Fig. 2C) indicating low rates of recombination, and possibly also some accumulation of new mutations. Many of the sexual populations in the Atlantic and the Transition zone also showed negative *F*_IS_ values (Fig. 2C). This corroborates our earlier observations of occasional clonemates in the NJ tree as negative values of *F*_IS_ reflect loss of recombination: values of *F*_IS_ between -0.05 and -0.15, observed among the populations outside the Baltic Sea, suggest regular asexual recruitment in these populations (42-44). Consequently, our data indicate that *F. vesiculosus* is currently facultatively asexual also outside the Baltic Sea, and that cloning was already present when the colonisation of the Baltic Sea began.

Among the clonal clusters, 34 and 49 were different from the other four clusters with higher genetic variation (*H*_e_), and, in particular 34 also had a value of *F*_IS_ close to zero. A demography involving more recombination events would explain such *F*_IS_ values, and as indicated by the NJ tree, these clusters also consisted of higher proportions of discrete clones than the other clonal clusters (Fig. 1A). Alternatively, the long branches represent old coalescent times of clones, and if so, the small clones inside the cluster would instead be part of the same clonal lineage that had diverged without involving recombination.

### Modelling the genetics of the colonisation

We explored the dynamic relationships between *H*_e_, *H*_o_, and *F*_IS_ in facultative asexual species using a one-dimensional model to represent an expansion along a coastline, such as in *F. vesiculosus* from the Atlantic and into the Baltic Sea. Simulations covered temporal patterns up to 10,000 reproductive seasons (=years) in *H*_e_, *H*_o_, and *F*_IS_ over a chain of 500 patches connected by symmetric gene flow. The first 50 demes represented the core meta-population (“in the Atlantic”) and were allowed to stabilise during a burn-in period of 250,000 reproductive seasons prior to expansion. Each deme had a maximum *N* of 100 individuals. The model was parameterised with values relevant for *F. vesiculosus*, most importantly, separate sexes and facultative asexual reproduction, larger per-individual investment in sexual than in asexual reproduction, dispersal only to nearby sites unless occasional long-distance migration occurred, and competition for space in each site. We included mutations in the model to illustrate their effect on genetic variation, and newly formed clones either remained sexually competent or successively lost their sexual capacity at a certain rate due to, for example, the accumulation of deleterious mutations (for model details see Methods).

Model outputs from single realisations allowed us to visualise the stochastic nature of the patterns displayed. For this purpose, we chose realisations with representative parameter sets that resulted in the establishment and persistence of one or more local clones during at least 500 reproductive seasons. In the model we assumed that demes with genotypic identity (based on individuals’ ID; see Methods) of at least 0.95 belonged to the same local clone. In addition, among these representative parameter sets, we gathered those where long-range dispersal was absent, and we ran a further 200 independent simulations for each such set to obtain the average patterns. Note that in the model with long-range dispersal, the timing and location of the established clonal colonies is expected to be highly variable between individual realisations, meaning that averaging over independent realisations would not be informative.

#### Model outcome with short-range dispersal

We first could confirm the findings of the earlier model (11) that a facultative asexual species with migration only between neighbouring demes colonise an empty one-dimensional landscape through a “clonal wave” formed at the front of the expansion, and thereafter a sexual wave followed (Fig. 3A-C). We already knew that unless sexual recruitment is relatively less successful than asexual recruitment this is a very robust prediction (11). Furthermore, under high rates of dispersal the sexual wave spread more efficiently than when dispersal was slow (Fig. 3D-F), and the spread of the sexual wave is faster the higher the success rate of sexual reproduction is relative to asexual reproduction (11). In our current model we also introduced successive loss of sex in clones, but this did not affect the spread of the clone under low rates of dispersal (Fig. 3C), while loss of sex prolonged the life of the clone under high rates of dispersal (Fig. 3F).

**Fig. 3.**
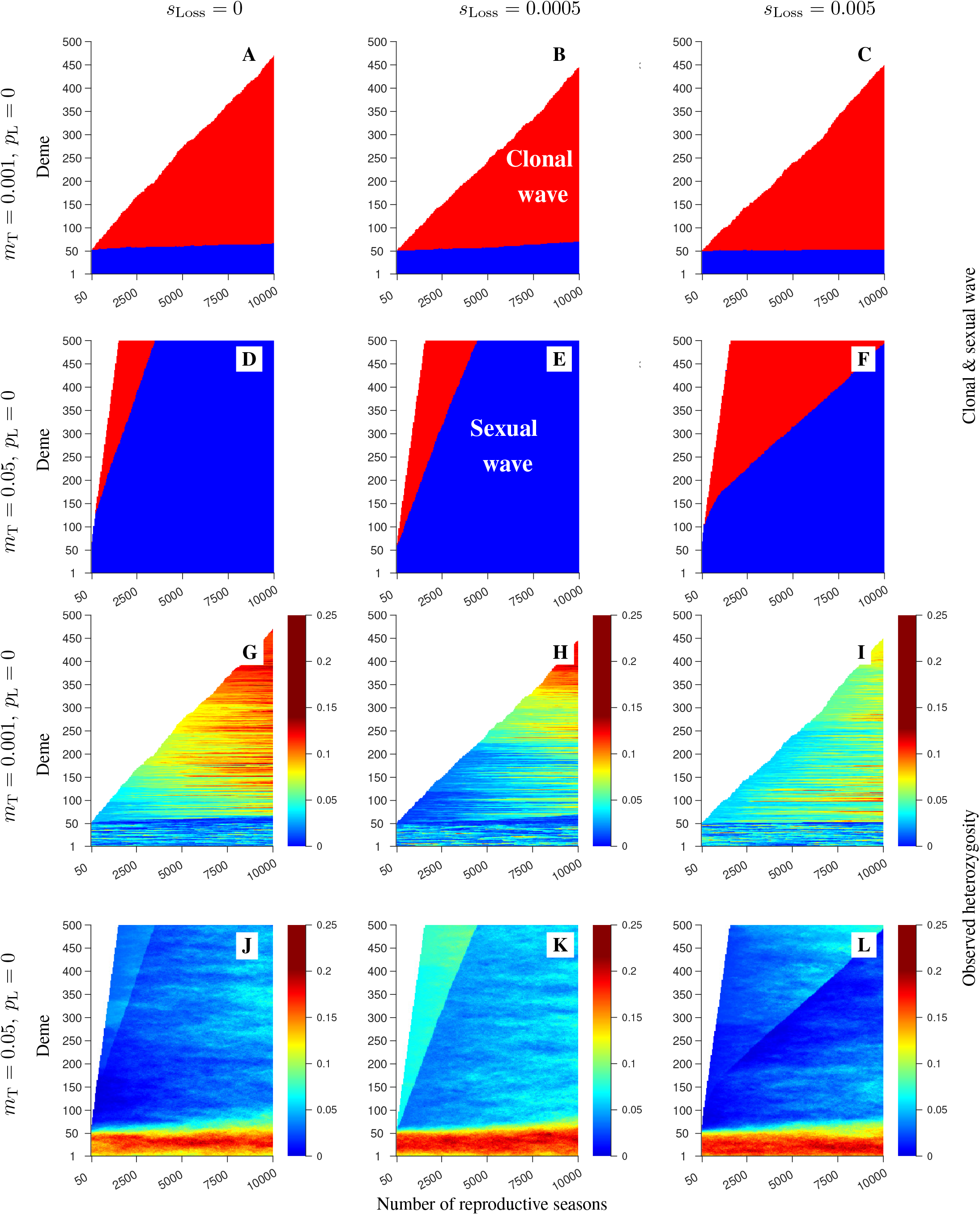
Simulation results from single realisations of the model where dispersal occurs only to the nearest neighbours. Panels in the upper two rows show the spatio-temporal dynamics of the pioneer clone (red) and sexual areas (with potentially rare clones; blue). Empty areas are coloured white. The rows differ by the migration rate: 0.001 (**A**-**C**), or 0.05 (**D**-**F**). The columns differ by the rate of loss of sex (*s*_Loss_) as indicated in the figure. For each panel in rows 1-2, the spatio-temporal dynamics of observed heterozygosity averaged over all loci (depicted by the colour map) are shown in the corresponding panel in row 3-4 (and see Suppl. Fig. S1 for *H*_e_ and *F*_IS_). Remaining parameters: total number of demes *K* = 500, maximum number of individuals in each deme *N* = 100, death rate *d* = 0.1, number of mating attempts per fully sexually active male *S* = 8, number of clonal offspring per inidivudal *C* = 1, number of loci *L* = 100, mutation rate *μ* = 0.000025, free recombination between any pair of loci, probability of long-range dispersal *p*_L_ = 0. The burn-in was run for 250,000 reproductive seasons wherein only demes 1,2, …, 50 were fully occupied and the remainder of the habitat was empty.

Our new model outlined the complex dynamics of *H*_o_, *H*_e_, and *F*_IS_ along the 500 demes and over a time period of 10,000 reproductive seasons (Fig. 3G-L showing *H*_o_, and Suppl. Fig. S2 showing *H*_e_ and *F*_IS_ for the same model realisations as in Fig. 3A-F). Under low dispersal (0.001, Fig. 3G-I), genetic drift rapidly eroded the genetic variation in core demes (1-50). Consequently, the clone emerging from the edge of the core population carried a low level of heterozygosity. Over time, however, the accumulation of new mutations made this clone increasingly heterozygote, and increasingly so the more distant from the core demes. Conversely, the sexual demes remained on a level of genetic variation set by the mutation-drift-migration balance, which for low rates of migration meant generally low expected and observed heterozygosity levels. Under high migration, by contrast, much higher levels of genetic variation were maintained in the core demes (Fig. 3J-L). Still, the emerging clonal wave carried a low proportion of heterozygote loci at take-off from the sexual core population. This was at first glance an unexpected outcome of the model, however also observed in the empirical data (i.e. low *H*_e_ in four of the clones compared to sexual populations, see Fig. 2), and we will return to this intriguing observation below after first considering the effect of long-range dispersal.

#### Adding occasional long-range dispersal

If long-range dispersal at a low rate was introduced into the system, the spatial structure changed dramatically (Fig. 4 and Suppl. Fig. S3). The initial wave of the pioneer clone was soon followed by the spread of clones to new demes and the formation of additional clones. Long-range dispersal introduced single individuals of different genotype and sex into demes where a clone (e.g. the pioneer clone) was already established. This resulted in sexual activity and formation of a new clone that occasionally spread and was able to establish in a new and empty area. Formation and establishment of new clones happened most readily under relatively low short-distance migration and less rare long-distance migration (Fig. 4A-C). Under more frequent short-distance migration and rarer long-distance migration fewer new clones were established, and instead the sexual wave was more prominent (Fig. 4D-F). If sex was not lost, the sexual wave assimilated the clones after some time (Fig. 4A, D), but if clones lost the capacity of sex relatively quickly (through the accumulation of deleterious mutations), they resisted recombination and remained intact for extensive periods of time (Fig. 4C, F). Empty demes in between clonal demes sometimes developed into pockets of sexual populations upon the arrival of several long-distance (or short-distance) migrants of different sex (Fig. 4C, dark blue horizontal stripes between clonal sections).

**Fig. 4.**
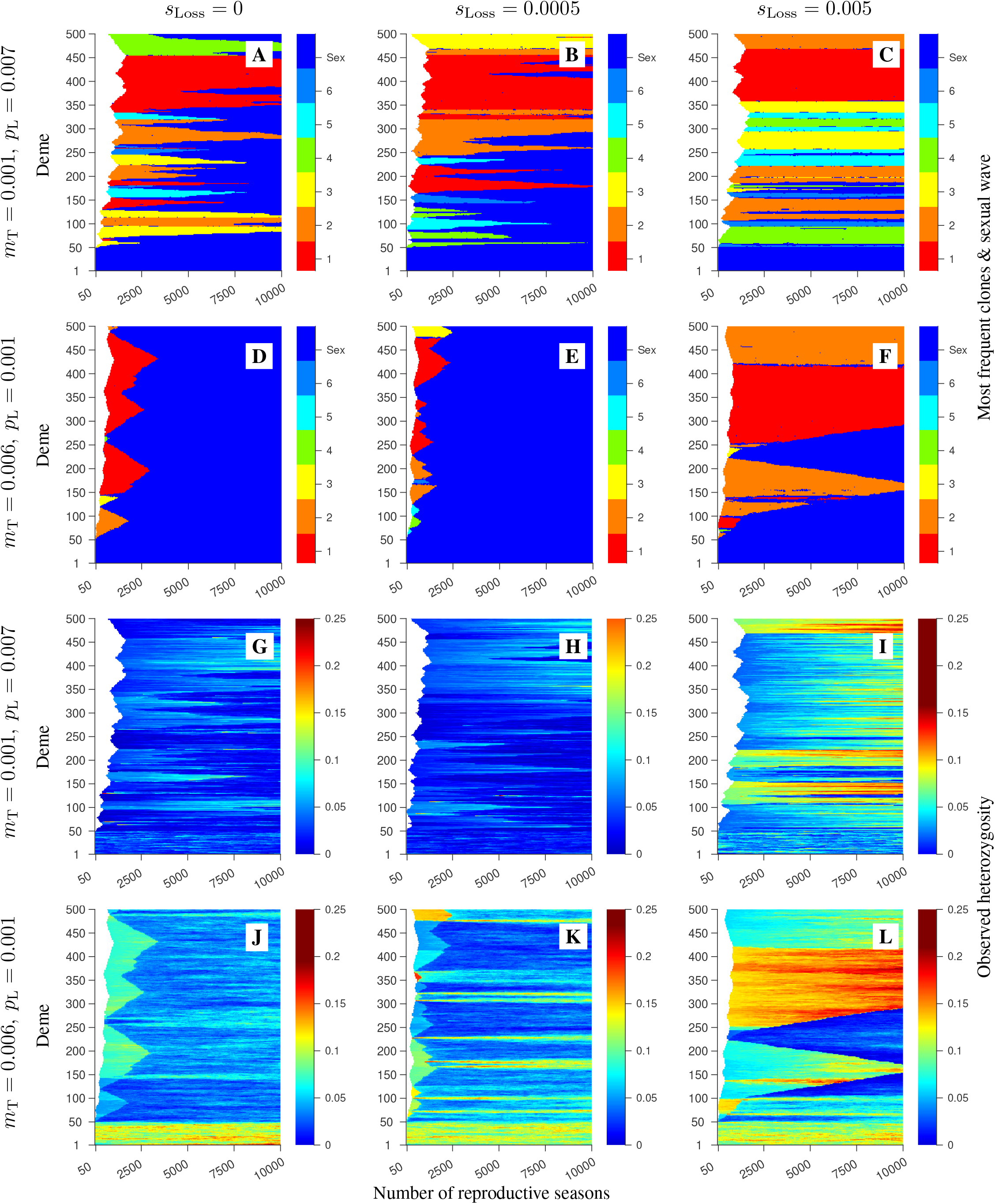
Simulation results from single realisations of the model with both short- and long-range dispersal. For each panel in rows 1-2, the spatio-temporal dynamics of observed heterozygosity averaged over all loci (depicted by the colour map) are shown in the corresponding panel in row 3-4 (and see Suppl. Fig. S2 for *H*_e_ and *F*_IS_). Captions are same as in Fig. 3. The rows differ by the total migration rate (*m*_T_) and the probability that a migrant disperses by long-range dispersal (*p*_L_) as depicted in the figure. The columns differ by the rate of loss of sex (*s*_Loss_) as indicated in the figure. The mean long-range dispersal distance was set to 50 demes. The remaining parameters are as in Fig. 3.

As before, heterozygosity in core demes was very sensitive to rates of short-distance migration (Fig. 4G-L), and under most scenarios variation was lost upon formation of clones. However, occasionally a dominant clone was formed with relatively high *H*_o_ (Fig. 4L, e.g. red clone in 4F has attained higher *H*_o_ than the orange clone), and as before, surviving clones also accumulated new variation from new mutations.

#### Patterns of genetic variation along the route of the expansion

To compare the model outcome with the empirical trajectories of *H*_o_, *H*_e_, and *F*_IS_ along the Atlantic - Baltic Sea transect, the spatial patterns of average values *H*_o_, *H*_e_, and *F*_IS_ generated from 100 model realisations of each model were plotted along the experimental demes 1000, 2000 and 10,000 reproductive seasons after the start of the expansion in the model with only short-range dispersal (Fig. 5 shows the result after 2000 generations, and see Suppl. Figs. S4 and S5 for 1,000 and 10,000 reproductive seasons). As already suggested from the single realisations of the same model (Fig. 3), low rates of short-range dispersal caused genetic variation to drop already in the core metapopulation, and this pattern was confirmed in the average plots (Fig. 5A-C, G-I). Both *H*_e_ and *H*_o_ dropped from 0.2 (the starting value in all simulations) to values below 0.05 in the core demes (1-50) already before the first clone was formed (Fig. 5; green bars overlaying blue bars illustrates variation in core demes at the end of the burn-in period). The drop in genetic variation and heterozygosity was even more pronounced at the edge of the core meta-population, from which the first clone was formed (Fig. 5A-C). The shift from dominance of sexual (blue) to clonal (red) individuals accompanied by a drop in *F*_IS_, took place only a few demes away from the core demes for low rates of dispersal this (Fig. 5M-O).

**Fig. 5.**
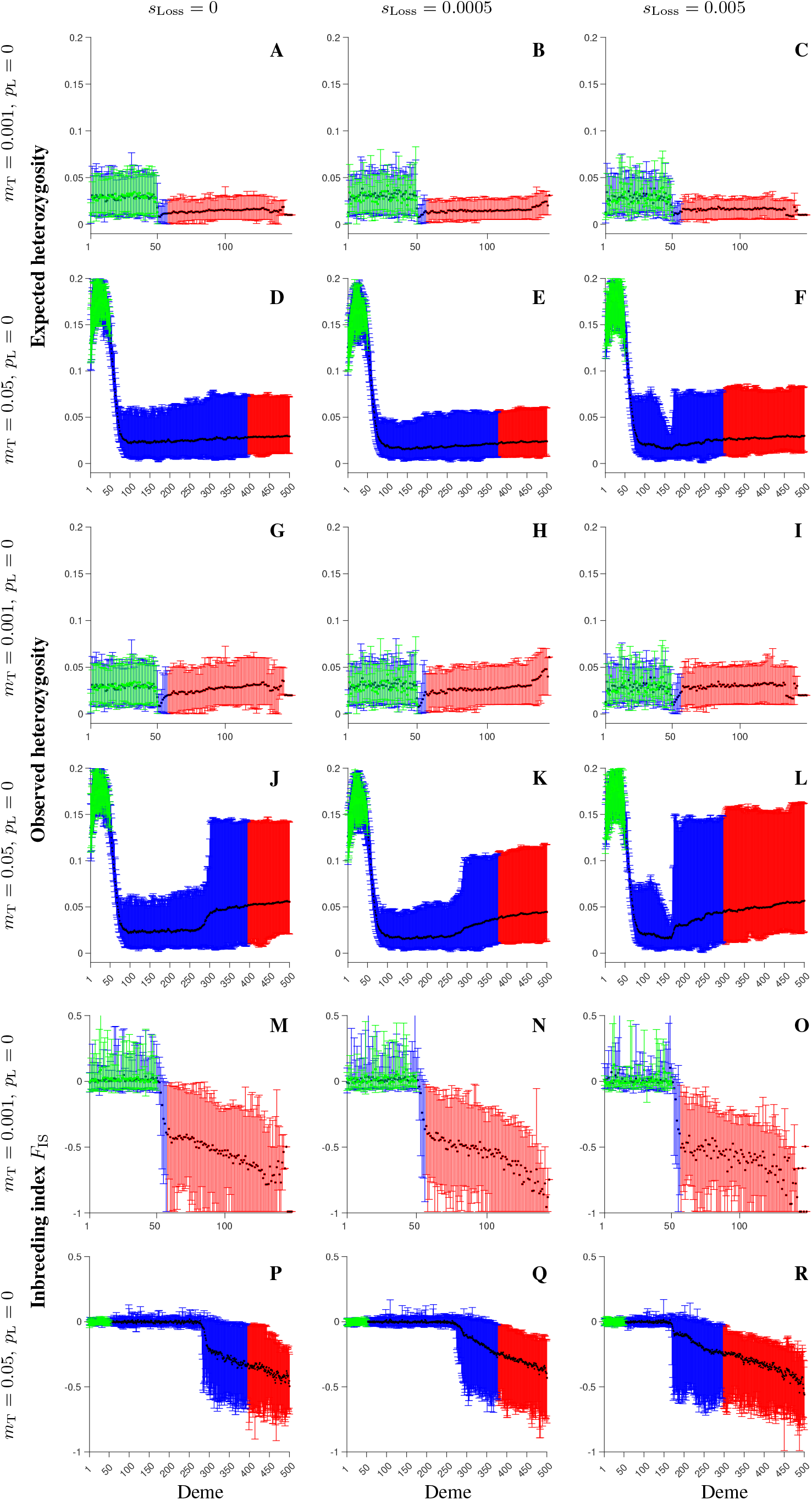
Population genetic structure with only short dispersal after 2000 reproductive seasons. Expected heterozygosity (**A**-**F**), observed heterozygosity (**G**-**L**), and *F*_IS_ (**M**-**R**) were obtained 2000 reproductive seasons after the start of expansion in the model where dispersal occurs only to the nearest neighbours. The black symbols show the values averaged over 100 independent realisations, the blue and red bars span between the 5^th^ and 95^th^ percentiles of the realised distributions in the sexual and clonal areas, respectively. For comparison, the green symbols and error bars show the corresponding results at the end of the burn-in simulations wherein only demes 1,2,…, 50 were fully occupied and the remainder of the habitat was empty. The rows differ by the value of the migration rate, and the columns differ by the rate of loss or sex, as depicted in the figure. The remaining parameters are as in Fig. 3.

Under higher rates of short-distance dispersal, genetic variation and heterozygosity were maintained among core demes with only a minor decline towards the edges of the core meta-population (Fig. 5D-F, J-L). The reason for this was that high dispersal among demes essentially compensated the loss of variation due to genetic drift in local demes. However, at the start of the expansion, both *H*_e_ and *H*_o_ dropped rapidly when the new demes were colonised during the expansion. The reason for this was a series of bottlenecks and loss of variation from drift during the initial phase of colonisation before a first successful clone was able to take-off and form a clonal wave. That is, the first empty deme to be colonised received migrants of both sex before it was completely filled up (due to the relatively high rate of short-distance dispersal), and this prevented the establishment of a completely clonal deme from which the next deme could be colonised. After a couple of such trials, a clone eventually filled up a new deme completely before the arrival of another sex, and a clonal wave was initiated. Hence, following repetitive bottlenecks, the first clone took off from a strongly depauperate sexual population. The model realisations show that the loss of genetic variation by drift leveled out quickly at around demes 70-80 (Fig. 5D-F, J-L), which suggests that around here the pioneer clone was formed and run away from the sexual population. After 2,000 reproductive seasons, the sexual wave (blue) had spread into the area where the clone first established, but less so if the clone rapidly lost sexual capacity (Fig. 5F, L, R). Adding long-range dispersal, patterns of individual realisation showed similar trends as for only short-range dispersal, with initially strong drop in genetic variation before one, and later additional clones, were formed, although the overall structures were highly complex (Suppl. Figs. S6-S8).

### Comparing model output with the empirical patterns

The model showed that with only local dispersal, a single large clone would dominate the asexual phase of the expansion and fill up the new territory, while with occasional long-range dispersal new clones would establish and potentially also spread to other areas. We found a handful of large clonal clusters at the margin of the Baltic Sea range, five with local distributions and one established along 600 km of coastline. The appearance of several clones suggested an interaction between sexual and clonal recruitment, such as initiated by occasional long-range dispersal in the model. Indeed, the complex and dynamic population genetic structure achieved in the model when adding occasional long-range dispersal resembled the complexity we observed in the empirical data, and from this we conclude that occasional long-range dispersal during establishment has been a key component shaping the spatial structure of *F. vesiculosus* in the Baltic Sea.

Our model also predicted a substantial loss of genetic variation in the sexual populations from which the pioneer clone was formed, resulting in the somewhat surprising finding of clones of low genetic variation. In fact, both the scenarios with low and with high rates of short-distance dispersal resulted in formation of clones from sexual populations that had already lost most of the original variation from genetic drift. Under low dispersal, this loss occurred already during the burn-in period and predominantly in the edge demes of the core meta-population, while under high dispersal, the core demes resisted genetic drift while the drop in variation occurred during the colonisation of the first demes along the expansion route. The genomic data clearly showed that four out of six clonal clusters had lost much genetic variation compared to all sexual populations, and so these clones must have been formed from sexual populations depauperate in genetic variation. We believe that the mechanism illustrated in the model realisations explains how this happened.

In the model, clones that remained sexually viable were assimilated into the sexual wave after some time, while a successive loss of sex prolonged the life-times of clones. While physical isolation protected a clone with sexual competence from recombining away during the expansion phase, the genomic data showed that currently several clones live side-by-side with sexual populations. In addition, the low levels of genetic variation of four of these clones also reflected their origin at the expansion front of the colonisation of the Baltic Sea and so that they were established before the arrival of sexual individuals. The sexual populations inside the Baltic Sea, on the other hand, all have a comparably high level of genetic variation similar to outside the Baltic Sea. Indeed, this was not predicted by any of the model scenarios; these instead showed that the sexual wave would also experience initial bottlenecks. We will return to this issue in the Discussion, but overall, consistency between model predictions and empirical patterns suggests that the processes and mechanisms outlined in the model scenarios can inform us about the evolutionary processes shaping the complex seascape of *F. vesiculosus* in the Baltic Sea.

## Discussion

Increased asexuality in marginal areas of a species’ distribution (“geographic parthenogenesis”) is frequently observed in plants, but also in some animal species (6, 8, 50-53). One hypothesis to explain these observations is the fixation of the most suitable genotype in the marginal environment and avoidance of migration load caused by suboptimal alleles arriving from more central populations (6, 7, 54). An alternative suggestion is that selection favours asexual reproduction in small and fragmented populations to avoid inbreeding depression (55). Here we further explore a third hypothesis that does not invoke selection among genotypes. The key mechanism is instead the uniparental advantage and reproductive assurance of clones during establishment in a new, and potentially marginal, area. This mechanism gives an advantage to asexual individuals over sexual individuals of dioecious or non-selfing hermaphrodite species that require two individuals of opposite sex to arrive simultaneously in the same site for sexual recruitment to take place (56-58). The asexual advantage in recruitment will result in a clonal wave that runs ahead of the establishment of sexual individuals and colonise new and/or marginal areas.

### Expansion of a facultative asexual species can involve a primary clonal and a secondary sexual wave

Using a mathematical model, we earlier showed that the reproductive assurance of clones will result in a clonal wave as a first step in colonisation of an empty habitat (11). We also showed that if asexual reproduction is less favourable in general than sexual reproduction (e.g. more costly), the clonal wave will after some time be followed by a sexual wave that eradicates the clone (11). To evaluate the model predictions of a primary clonal wave followed by a sexual wave, we generated extensive genomic data for the seaweed *Fucus vesiculosus*, earlier considered obligate sexual but since the discovery of clones inside the Baltic Sea (36) turn out to be a facultative asexual species. However, before addressing the hypothesis of formation of a primary clonal and a secondary sexual wave during colonisation of a new habitat, we needed to understand the origin of asexual reproduction by asking two questions: First, where did asexual reproduction evolve? Was cloning already present outside the Baltic Sea or did it arise as a new trait inside this young environment, as earlier suggested (36)? Second, did clones have multiple origins as predicted from occasional long-range dispersal in our earlier model (11), or were they all from the same lineage?

With extensive genomic data we are now able to answer the first question by showing that asexual reproduction is also present outside the Baltic Sea and cloning is an ancestral trait to the Baltic lineage of *F. vesiculosus*. It is already well-known that this species forms adventitious branches in both brackish and marine waters (59), but it has earlier been suggested that this was part of a wound-healing mechanism (60). Asexually recruited individuals are morphologically indistinguishable from sexually recruited individuals make clones only detectable using genetic markers, hence the occurrence of clones has been overlooked by biologists for centuries. The finding of asexual reproduction being an ancestral trait has two implications: First, there is no need to invoke the rather unlikely scenario that a completely new reproductive trait evolved immediately upon colonisation of the Baltic Sea only 8,000 years ago. Second, a geographic parthenogenesis-like pattern in *F. vesiculosus* is an expected outcome of facultative clonal reproduction in combination with slow stepping-stone migration (11, this study), and not a spurious effect of a rare mutation inside the Baltic Se, and we should therefore expect that under similar conditions elsewhere in *F. vesiculosus*, and in other species, clonal waves can spread and establish in recently occupied areas.

Our genomic data also answered the second question, showing multiple origins of clones. As predicted from our earlier model, adding occasional long-range dispersal to a system dominated by short-range dispersal would result in clones being formed repeatedly from different recombination backgrounds (11). That short-range dispersal is the predominant dispersal mechanism is obvious from both our genomic data and earlier genetic studies clearly showing a small-scale spatial structure. For example, Tatarenkov et al. (36) found that genetic divergence at four spatial levels (10 m, 1 km, 70 km and 700 km) contributed roughly equal amounts to the total divergence. However, occasional long-distance dispersal may occur by drifting pieces of thalli which remain sexually active if they contain mature gametes (30), or more likely, remain asexually active by shedding adventitious branches upon arrival at a new site.

### Extensive genomic data and model scenarios explain unexpectedly low genetic variation in clones

Knowing that cloning is an ancestral trait in *F. vesiculosus*, we used our empirical data to test the main prediction deriving from our earlier model, the occurrence of a primary clonal wave and a secondary sexual wave during colonisation of the Baltic Sea by *F. vesiculosus*. To do this, we used our new model to elaborate the spatial genetic structure of the expansion by analysing the dynamics of the population genetic descriptors *H*_e_, *H*_o_ and *F*_IS_. This was a necessary step to be able to better understand the mechanisms and processes giving rise to a clonal and a sexual wave. For example, a key prediction from the new model was a decreased genetic variation in the clonal wave compared to the sexual wave, and our empirical data supported this in four out of six clonal clusters. The loss of genetic variation in the clones supported formation of clones early in the colonisation process from depauperate sexual populations at the expansion front, rather than from the sexual populations currently established in the Baltic Sea. Hence, this is a key result supporting the formation of clones at the initial stage of the colonisation, with sexual populations now side-by-side with the clones in some areas, arriving at a later stage.

On the other hand, none of our model realisations indicated increased genetic variation during or following the sexual wave, as observed in the empirical data. In our model, deme size was rather small (N=100) and migration was one-dimensional and restricted (0.001-0.05), and hence genetic drift was predominant in maintaining a low level of genetic variation in all demes, except in the core demes and close to these under the higher rate of migration (e.g. Fig. 3J-L). A tentative explanation for the high genetic variation observed in the current sexual populations in the empirical data is that over time deme sizes increased, as did the total metapopulation size in the Baltic Sea, reducing the effect of drift. Currently, *F. vesiculosus* is a dominant and widespread perennial seaweed inside the Baltic Sea, and it occupies most available rocky intertidal and shallow subtidal habitats throughout its area of distribution. However, such a successful establishment might not have happened immediately, but rather slowly after an initial period of high selective mortality and low effective populations size. Current sexual populations are locally adapted in several different traits (61-63), and this might have paved the way for larger effective population sizes and increased genetic variation mediated by gene flow from outside the Baltic Sea. Today the Baltic Sea meta-population of *F. vesiculosus* is genetically diverged from the Atlantic (Fig. 1A), and the peak in genetic variation in the Transition zone (Fig. 2A) maybe suggests that migration mix two different gene pools in this area.

### Large clones can be persistent in marginal areas

In our original model (11) the sexual wave always assimilated the clonal lineages unless asexual reproduction was less costly than sex and impeded the spread of a sexual wave. Accumulation of deleterious mutations that hamper the intricate sexual mechanisms has been observed in clonal species (8, 64), and is expected because such mutations will appear neutral during asexual expansion. As we show in our new model, if the capacity for sexual reproduction is successively lost in a clone, this will prolong survival extensively of those clones that escaped during the early phase of the expansion, while younger clones will be caught up by the sexual wave while they are still sexually competent. For those clones that escape the sexual population and managed to spread ahead of the sexual wave, the loss of sexual competence will make them able to survive side-by-side with sexual individuals, as we observe in the Baltic *F. vesiculosus*. Possibly, a negative environmental impact on gamete function in *F. vesiculosus*, such as polyspermy described from low salinity areas in the Baltic Sea (65, 66) might further support clonal survival in the marginal areas. However, sexual populations are currently established in low salinities (40, this study) hence salinity cannot be a major determinant of the rate of sex. Consequently, clones of different sex have also been found mixed at a scale of meters without signs of recombination (38), suggesting that these clones are now more or less completely asexual.

Despite *F. vesiculosus* also being facultatively asexual outside the Baltic Sea, we did not find large and dominant clones in the Atlantic and the Transition zone. Formation of new clones is likely to take place at all sites through dropping of adventitious branches. However, in an established dense sexual population re-assimilation of new clones into the sexual population will start immediately, if the asexually recruited individual remains sexually competent. Unless formation of adventitious branches is much less costly than formation of gametes, in which case a species will soon be dominated by clones everywhere (11). Thus, the absence of large clones of *F. vesiculosus* in the Atlantic meta-population supports our model assumption that asexual recruitment is generally less successful than sexual recruitment, except at the front of an expansion into an empty habitat.

### Spread of a facultative asexual species can result in a complex and dynamic population genetic structure

According to the models, the formation and spread of a pioneer clone, additional clones, and finally a sexual wave, result in a complex genetic landscape. This is also what we observe in *F. vesiculosus* inside the Baltic Sea (38, 40, this study). The resulting mix of genetically homogeneous clones and small-scale heterogeneous sexual populations results in a patchy genetic structure where the level of genetic variation is difficult to predict at any given site. Moreover, the structure is likely to be transient in time, at least on small spatial scales (<1km) (38). This can be due to short-term stochastic fluctuations or long-term trends of gradual replacement of clones by sexually recruiting individuals with capacity to evolve new adaptations to a changing environment. Additionally, clones are expected to successively degrade from increasing loads of deleterious mutations (58, 67) and unless somatic mutations provide the necessary genetic variation for selection among clones of a clonal cluster, leading to adaptation to a changing environment (21), clones will decay over time. The extensive distribution of a large female clone over 600 km of coastline, encompassing gradients of both salinity and temperature, is intriguing, and somehow suggests that adaptation within a clonal lineage might be possible, although extended empirical work is certainly needed to evaluate such a suggestion.

As we show here, the genetic complexity generated by the clonal and sexual waves during expansion processes can be persistent at least over tens of thousands of generations and will set the stage for a species to potentially adapt to environmental changes. Under the ongoing global challenges of a changing climate, not least affecting enclosed marine basins like the Baltic Sea, management authorities are challenged by the need for addressing the genetic diversity of species and how to protect such diversity. For this, a very intense sampling and detailed plans for protection and conservation might be needed, if, as in the Baltic *F. vesiculosus* populations, a sexual population in one area might be replaced by a single clone just a few kilometres away.

## Materials and Methods

### DNA Extraction and Genotyping

We extracted genomic DNA of 20 individuals per locality from 47 localities (Suppl. Table 1) and re-extracted 4 of those 20 individuals from each sampling location to use as technical replicates (i.e. replicated extraction, library preparation and sequencing). We followed Panova et al. (68) for genomic DNA extractions, grinding dried algal tissue and washing it twice with 100% acetone before digestion in CTAB buffer. Then, we extracted the DNA using the DNA Plant II Extraction Kit (Qiagen) with an extra cleaning step using the DNA Clean and Concentrator kit -25 (Zymo). We assessed the DNA integrity by electrophoresis in a 1% agarose gel and the quality using Nanodrop. We quantified the DNA using a Qubit dsDNA broad range AssayKit (Invitrogen-ThermoFisher Scientific). We constructed individually barcoded 2b-RAD libraries (41) following a protocol modified by Aglyamova and Matz (https://github.com/z0on/2bRAD_GATK/blob/master/2bRAD_protocol_june1_2018.pdf) as described in Kinnby et al. (63).

We used the computer cluster ‘Albiorix’ at the University of Gothenburg, Sweden, for all our bioinformatic analysis, including a modified de novo pipeline for trimming, filtering and genotyping, available at https://github.com/crustaceana/TheFucusProject and originally written by Mikhail Matz (https://github.com/z0on/2bRAD_denovo/blob/master/2bRAD_README.sh). Sequences were trimmed and quality filtered before being mapped to the draft genome of *F. vesiculosus* (63). We subsequently called the genotypes with the GATK pipeline and used the technical replicates to extract reproducible SNPs across all replicates and for non-parametric quantile-based recalibration of variants. We removed loci exceeding 75% heterozygotes that may be a result of sequencing errors and filtered individuals with excess false homozygotes due to poor coverage or excess false heterozygotes based on per-individual inbreeding coefficient. After variant recalibration, we removed the technical replicates and thinned the dataset to retain one SNP per RAD fragment. We performed further thinning to remove loci or individuals with >5% missing data and to retain only informative loci and to avoid loci potentially affected by null alleles using the R package “radiator” (69). All raw 2b-RAD sequences are deposited in the Sequence Read Archive repository at the National Centre for Biotechnology and Information (NCBI BioProject Number PRJNA629489).

### Evolution of clonal expansion and detection of lineage-specific diagnostic loci

We first determined the number of clones in our dataset and the geographic patterns of clonal expansion with a complete dataset comprising 964 individuals distributed across 47 localities spanning France, UK, Norway and along the Baltic Sea in Denmark, Germany, Sweden, Finland, Latvia, Estonia, and Russia (Suppl. Table 1). We initially calculated genetic distances between individuals using the function diss.dist in the R package “Poppr” (70), which uses Prevosti’s genetic distance (71) to estimate the fraction of different SNP sites between samples. We used this genetic distance matrix to plot a Neighbour-joining (NJ) tree to identify the geographic distribution of clones and clonal clusters. We then calculated a conservative maximum genetic distance as cut-off for assigning clones using the ‘cutoff_predictor’ function in Poppr. The ‘cutoff_predictor’ function will filter out genotypes by degree of similarity using three clustering algorithms (Nearest Neighbour, UPGMA and Farthest Neighbour) to search the top fraction of genetic differences and find the largest difference. The average between the distances spanning that largest genetic distance is the cutoff threshold defining the clonal lineage threshold (70). After estimating the cut-off threshold, we identified similar multi-locus genotypes (MLGs or clones) using the mlg.filter function from Poppr to plot the frequency distribution of distances between samples in relation to the estimated threshold (Suppl. Fig. S9). The rationale for using this method to detect clonality is that if the distribution of genetic distances among samples shows high peaks toward low distances instead of a strict unimodal distribution, these peaks are likely to reveal somatic mutations resulting in low distances among slightly distinct MLG actually deriving from a single reproductive event (46-48, 72).

We further corroborated the presence of clonality on the identified MLG lineages by testing for linkage disequilibrium using the standardised index of association (rd) (73, 74) and compared them with two sexual populations outside the Baltic Sea and two within the Baltic (Suppl. Fig 10). Random sampling of the genome is expected to have a mean index of association close to zero for panmictic populations as opposed to populations under clonal reproduction (70). However, genome-wide sampling of potentially thousands of anonymous markers will likely create deviations from zero (random mating) even within sexual populations. Nonetheless, these deviations are expected to be considerably lower in sexual populations relative to clones. Following, the NJ tree revealed separate clonal clusters within the “Baltic” clade, suggesting the repeated independent evolution of clonal lineages. However, the tree cannot reveal reticulations among these clusters that shed light into the inter- and intraclonal contributions. We plotted a Minimum Spanning Network (MSN) with “Poppr” using our genetic distance matrix to reveal inter- and intraclonal reticulations and discern if the evolution of the clonal lineages has followed a single stepwise mode or if these lineages have evolved independently from each other. We furthermore characterised the early genomic signatures of genetic variation and clonality by calculating the relative levels of observed (*H*_o_) and expected (*H*_e_) heterozygosity, and inbreeding coefficient (*F*_IS_) for each lineage per locality using the R package “radiator”. Please note that the estimates of *H*_o_ and *H*_e_ are not showing absolute levels, because the RAD dataset has been filtered for polymorphic sites and with a minimum threshold of minor allele frequency.

### Spatial and temporal modelling of clonal expansion

Based on our experiences from the model of Rafajlović et al. (11) we constructed a new model and explored several different scenarios of expansion into an empty territory in a facultative asexual species. We focused on the temporal and spatial patterns of genotypic identity among individuals (based on their IDs; see below), expected heterozygosity, observed heterozygosity, and the inbreeding index *F*_*Is*_. We assumed a dioecious population, with diploid individuals capable of reproducing both sexually and asexually. The habitat consisted of *K =* 500 equally sized demes arranged linearly with equal distance between pairs of consecutive demes. We refer to the distance between pairs of consecutive demes as a unit distance in the habitat. In the simulations, we set the maximum number of individuals (N) in any deme to N = 100. The lifecycle of individuals consisted of: Migration (mostly to nearby neighbouring demes, but in some simulations we also allowed for long-range migration), production of offspring locally in each deme (sexual and asexual, with recombination preceding the sexual component of reproduction, plus mutation in either case), death of adults (assuming perennial populations with a per individual, per reproductive season death rate *d*), and local population-size regulation.

Note that each simulation was initialised in such a way that only the first *K*_0_ = 50 demes (i.e. demes numbered 1,2, …, 50) were occupied (hereafter referred as the source demes representing the core population), and the remaining demes were empty. We first ran a burn-in period accounting only for the source demes to allow the source population to stabilise under drift, migration and mutation. The duration of the burn-in was chosen to be much longer than the expected time to the most recent common ancestor in a well-mixed randomly mating population with *NK*_0_ individuals, with *d*^−1^ reproductive seasons during individuals’ lifetime, and it was set to 5*Nd*^−1^*K*_0_ (this corresponded to 250,000 reproductive seasons for *d* = 0.1. After the burn-in, we simulated an additional 10,000 reproductive seasons during which the population was allowed to expand over the empty demes.

In each simulation, migration from each deme occurred with the per individual, per reproductive season probability of *m*_T_. We assumed that an individual that migrates does so either by long-range dispersal (with probability *p*_L_), or by short-range dispersal (probability 1 − *p*_L_). In cases with *p*_L_ = 0, only short-range dispersal occurs. Otherwise, for 0 < *p*_L_ < 1, both types of dispersal occur (we did not model cases with *p*_L_ = 1, where only long-range dispersal would occur). We modelled short-range dispersal by assuming that an individual migrates either to its nearest left or right deme (if available), both demes being equally likely. Conversely, we modelled long-range dispersal by assuming than an individual migrates either to the left or right deme (both being equally likely) by a random distance sampled from an exponential distribution with mean *δ* assumed to be much larger than the unit distance, but much smaller than the total number of demes in the habitat, and we set *δ* = 50 throughout. Because we modelled a discrete habitat space, we discretised the exponential distribution of long-range dispersal distances similarly as in Eriksson and Rafajlović (9): Namely, the probability that an individual undergoing long-range migration would disperse by distance *D* was obtained by integrating the exponential distribution from *D* − 0.5 to *D* + 0.5. To separate long-range dispersal from short-range dispersal, we truncated this discretised distribution at distance 2 from below. For simplicity, we further truncated it at distance *K* from above. This was a justified practical simplification, because even prior to the truncation from the above, the probability of sampling a dispersal distance that is larger than or equal to *K* was negligible for the parameter values we used.

In either short- or long-range dispersal event, we did not allow individuals to migrate away from the habitat boundaries such that all individuals that would migrate to the left of deme 1 were placed in deme 1, and all individuals that would migrate to the right of deme *K* (or deme *K*_0_ during the burn-in) were placed at deme *K* (or deme *K*_0_ during the burn-in). In our simulations, all individuals were initially capable of reproducing both sexually and clonally, but clones could lose their sexual capacity over time (see below). We note that, due to the assumption of a maximal local population size, immigrants could only inhabit a deme they arrived in if there was space available (after the local emigration). Thus, if the number of local individuals that remained after emigration in a given reproductive season (*N*′; note that indexes for deme number and reproductive season are omitted here for simplicity) plus the number of immigrants was larger than *N*, we chose uniformly at random *N* − *N* ′ immigrants to inhabit the deme, whereas the remaining immigrants died. Otherwise, all immigrants were allowed in the deme.

For sexual reproduction, we assumed that each fully sexually active male (i.e. with sexual activity *s*_m,a_ = 1) was capable of encountering *S* individuals in its deme (via dispersal of sperm), where an encounter with a female that has not lost its sexual activity (i.e. that has *s*_f,a_ > 0) was assumed to result in one sexual offspring. Otherwise, for males with *s*_m,a_ < 1, the number of encounters was binomially distributed with *S* being the number of trials, and *s*_m,a_ the probability of success, and so the expected number of encounters was *s*_m,a_*S*. Note that we assumed that each female that has not lost its sexual activity completely (i.e. that has *s*_f,a_ > 0) is capable of producing as many sexual offspring as the number of encounters she experiences with sexually active males. This is a good approximation in situations when the number of sperms involved in each encounter is so large that one successful fertilisation is almost certain. Each encounter by a given male was assumed to occur with an individual chosen uniformly at random among all individuals in the male’s deme. Thus, the probability that an encounter will result in mating (and, thus, fertilisation) was 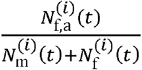, where 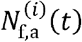 denotes the number of sexually active females with *s*_f,a_ > 0 in deme *i* in reproductive season *t*, 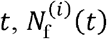 denotes the total number of females in deme *i* in reproductive season *t*, and 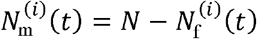 denotes the number of males in deme *i* in reproductive season *t*. Here, index *i* = 1,2, …, *K* denotes the deme number.

We traced the evolution of genetic variation in each deme simulating the allelic content at *L* = 100 loci, that were assumed to be bi-allelic (the two allele types were denoted 0 and 1), and to recombine freely (in an event of sexual reproduction). Mutations at each locus occurred according to a symmetric two-allele model, with a per allele, per offspring, per reproductive season probability *μ* fixed throughout each simulation run. In addition, given that all individuals were also capable of reproducing clonally, and it was necessary to trace the number of clonal lineages in each deme, we assigned an ID number to each individual and each sexually reproduced offspring received a new ID. Note also that we assumed that sexually produced offspring were either males or females with equal probability, and that all sexually produced male and female offspring had *s*_m,a_ *= s*_f,a_ = 1.

We modelled clonal reproduction in a given reproductive season such that each individual produced exactly *C* clonal offspring. We refer to *C* as the rate of clonal reproduction. Each clonal offspring inherited the sex and ID of its parent, as well as the full parental genotype prior to potential mutations. However, although the ID of any clonal offspring and its parent were identical, differences in the genotypes of a clonal offspring and its parent could occur due to mutations that were applied in the same way as for the sexual offspring (see above). In order to obtain clear information about the relatedness of individuals in each deme (especially where clonality was dominant), we used IDs of individuals to compute the genotypic identity. Note that if all individuals in a given deme are descendants of the same clone, the genotypic identity based on IDs would be equal to 1, whereas the genotypic identity based on genotypes that account for mutations would be less than 1. In addition, each clonal offspring was assumed to have a sexual capacity either equal to the sexual capacity of its parent (*s*_m,a_ for males, and *s*_f,a_ for females), or reduced in comparison to its parent by a non-negative constant denoted by *s*_Loss_. If the parent of a given clonal offspring lost its sexual activity (i.e., it had *s*_m,a_ = 0 or *s*_f,a_ = 0, depending on the sex of the parent), the offspring retained the value *s*_m,a_ = 0 (*s*_f,a_ = 0), independently of the value of *s*_Loss_.

Due to the competition for space, the generated offspring may or may not have survived, as explained next.

In the model we assumed that, after reproduction, any adult individual could die with the per individual, per reproductive season probability *d* (death rate), meaning that on average, each adult individual experienced *d*^−1^ reproductive seasons during its lifetime. Following the death of adults, we computed the number of living adults in a given deme and a given reproductive season, as well as the number of available places (*N*″; indexes for deme number and reproductive season are omitted here for simplicity) for offspring to occupy. This was done by subtracting the number of living adults from *N*. If the number of offspring produced in a deme was larger than the number of available places, we sampled uniformly at random *N*″offspring to survive from the list of offspring produced in this deme in a given reproductive season, whereby we accounted for both sexually and asexually produced offspring. Otherwise, all offspring survived.

Finally, we initialised the simulations (where only the source demes were inhabited) by setting the expected heterozygosity at each locus close to the value of the average expected heterozygosity estimated in the empirical data, i.e. 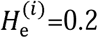 for *i* = 1,2, … *K*_0_, assuming (initially) Hardy-Weinberg and linkage equilibrium. But the source population was allowed to evolve thereafter during a long burn-in period, meaning that these initial assumptions are not restrictive. To generate the initial genotypes, for each locus, and each allele at the locus (recall that individuals were diploid) we chose uniformly at random among the alleles 0 and 1 with probability 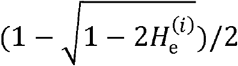 and 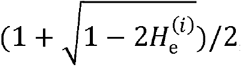, respectively. This was repeated independently for all loci and all individuals in the source demes.

In each simulation, the number of different clones in each deme (based on individuals’ IDs), genotypic identity (probability that a randomly chosen pair of individuals have the same ID), as well as heterozygosity (observed and expected) and *F*_IS_ at each locus were recorded every 50 reproductive seasons. Model parameters and values used in modelling is summarised in Table 1.

### Statistical analysis of population genetic estimators

We performed statistical tests for overall differences in genetic variation (*H*_e_) and observed heterozygosity (*H*_o_) among the three geographic areas, NE Atlantic, Transition zone and Baltic Sea both including and excluding the clonal lineages using the GenoDive 3.05 software (75). This included permutation tests to test for differences among geographic groups. OSx-statistic (76) was used as a test statistic comparing the sum of the squared differences over all pairwise combinations of groups. Permutations took place by randomising the populations over the groups.

## Acknowledgments

We greatly acknowledge help with field sampling around the Baltic Sea from Jonne Kotta, Merli Rätsep, Veijo Jormalainen, Luca Rugiu, Britta Meyer, Melanie J. Heckwolf, Carl André and Marina Panova, and for topical discussions with these and the rest of the BONUS-Bambi team. Finally, we are very grateful for useful comments on an earlier version of the manuscript from Martin Eriksson, Roger Butlin and Matteo Tomasini.

## Funding

Sequencing was performed by the SNP&SEQ Technology Platform in Uppsala. The facility is part of the National Genomics Infrastructure (NGI) Sweden and Science for Life Laboratory. The SNP&SEQ Platform is also supported by the Swedish Research Council and the Knut and Alice Wallenberg Foundation.

Swedish National Infrastructure for Computing (SNIC) at the High Performance Computing Center North (HPC2N, project SNIC 2021-5-501 provided computing time partly funded by Swedish Research Council VR, grant 2018-05973.

Swedish Research Councils VR and Formas, Linnaeus grant 217-2008-1719 (KJ main PI) EU and Swedish Research Council Formas, BONUS-Bambi project grant 2012-76 (KJ main PI)

Swedish Research Council Formas, grant 2019/00882 (KJ, MR)

2015–2016 BiodivERsa COFUND call (project MarFor), with national funding from Formas, grant 2016-01930 (KJ)

## Author contributions

Conceptualization: RTP, MR, KJ

Methodology: RTP, MR, MP, MT, PDW, KJ

Investigation: RTP, MR, PDW, AK, MP, MT, KJ

Visualization: RTP, MR, KJ

Writing—original draft: RTP, MR, KJ

Writing—review & editing: RTP, MR, PDW, AK, MP, MT, KJ

## Competing interests

Authors declare that they have no competing interests.

## Data and materials availability

Computer codes used for model simulations will be deposited to Dryad upon the acceptance of the manuscript. All the scripts used for the genomic analysis are deposited in Github (https://github.com/crustaceana/TheFucusProject/tree/master/scripts) and the genomic data into the Sequence Read Archive (SRA) from the National Centre for Biotechnology Information (NCBI) (BioProject Accession Number PRJNA629489).

## Notes

### Competing Interest Statement

The authors have declared no competing interest.

