## Supplementary Figs S1 to S10 and Supplementary Tables S1-S2 for "Clones on the run - the genomics of a recently expanded facultative asexual species"

**This PDF file includes:**

Figs. S1 to S10
Tables S1 to S2

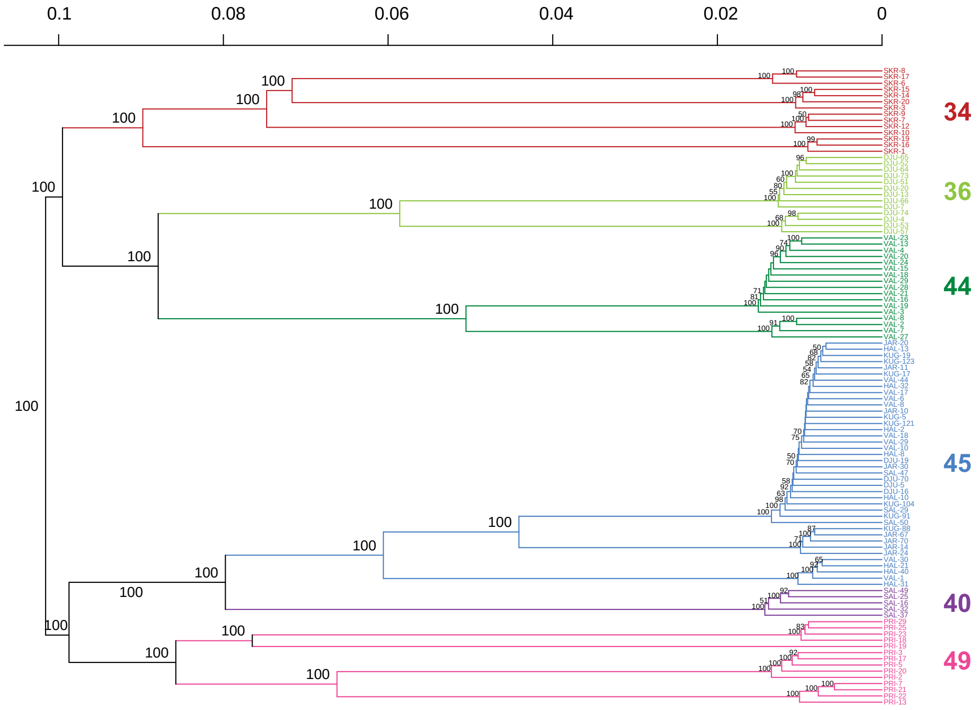

**Fig. S1.** Neighbour-joining tree <u>exclusively</u> of clonal clusters using K-means clustering through Bayesian Information Criterion in combination with bootstrapping, to determine the genetic delineation of each clonal cluster depicted with different colours. Numbers in each node are values after 1000 bootstraps. Large coloured numbers for each K lineage are according to Suppl. Table 1. Each individual is tagged with short site name (see Suppl Table S1) and ID number. The tree is calculated with a data subset and filtered separately for the purpose of showing the geographic affiliation of each individual within the clonal clusters, particularly of clone cluster 45. For this reason, the number of individuals on each clonal cluster may differ from Fig. 1 in main text, due to filtered individuals for missing data in the subset, but their genetic affinities have remained the same as in Fig. 1.

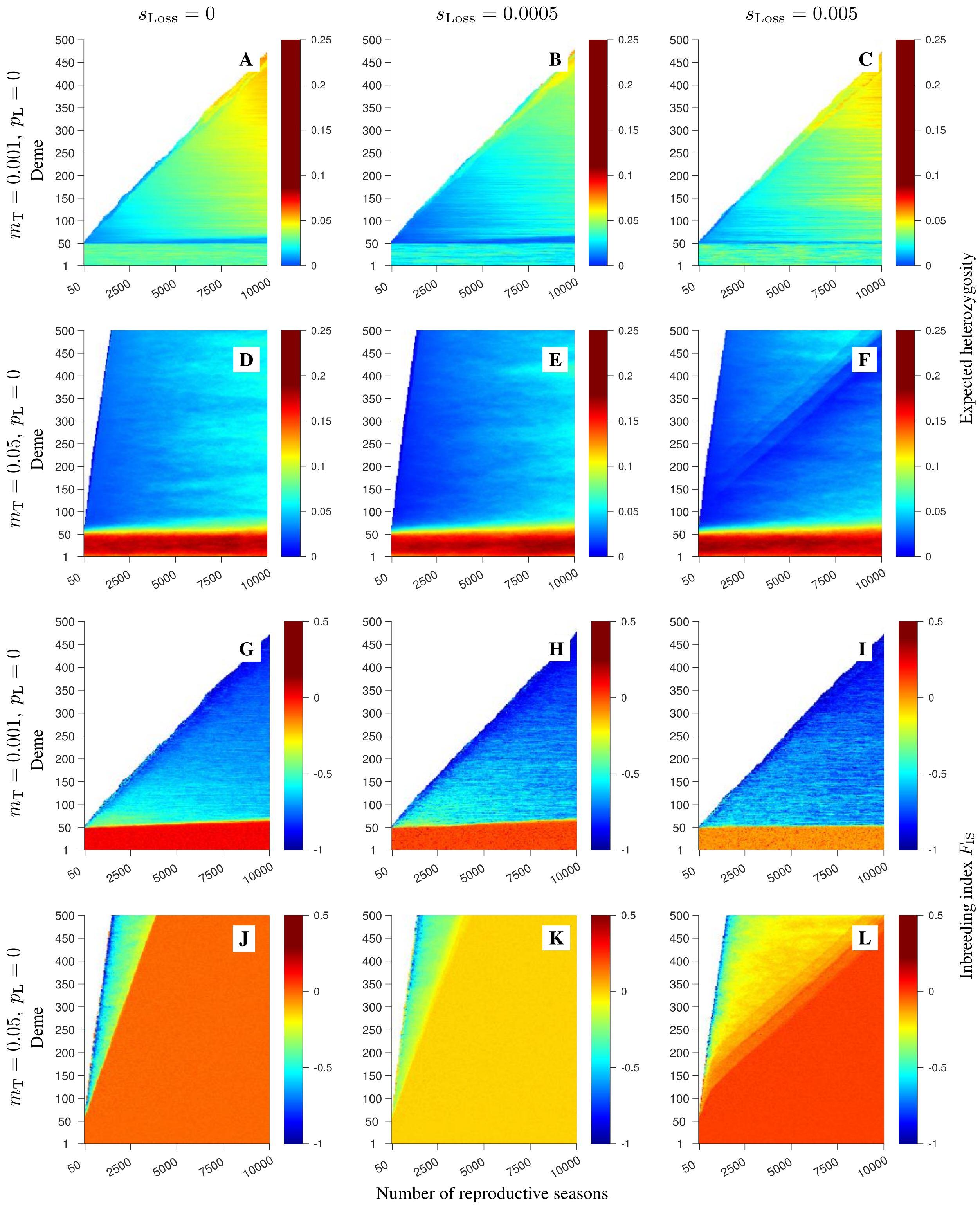

**Fig. S2**. Spatio-temporal patterns of expected heterozygosity and $F_{\mathrm{IS}}$ obtained in single stochastic realisations of the model with only short-range dispersal. The corresponding clonal structure and observed heterozygosity are shown in Fig. 3 in the main text. For the explanation of the parameters used, refer to Fig. 3 in the main text.

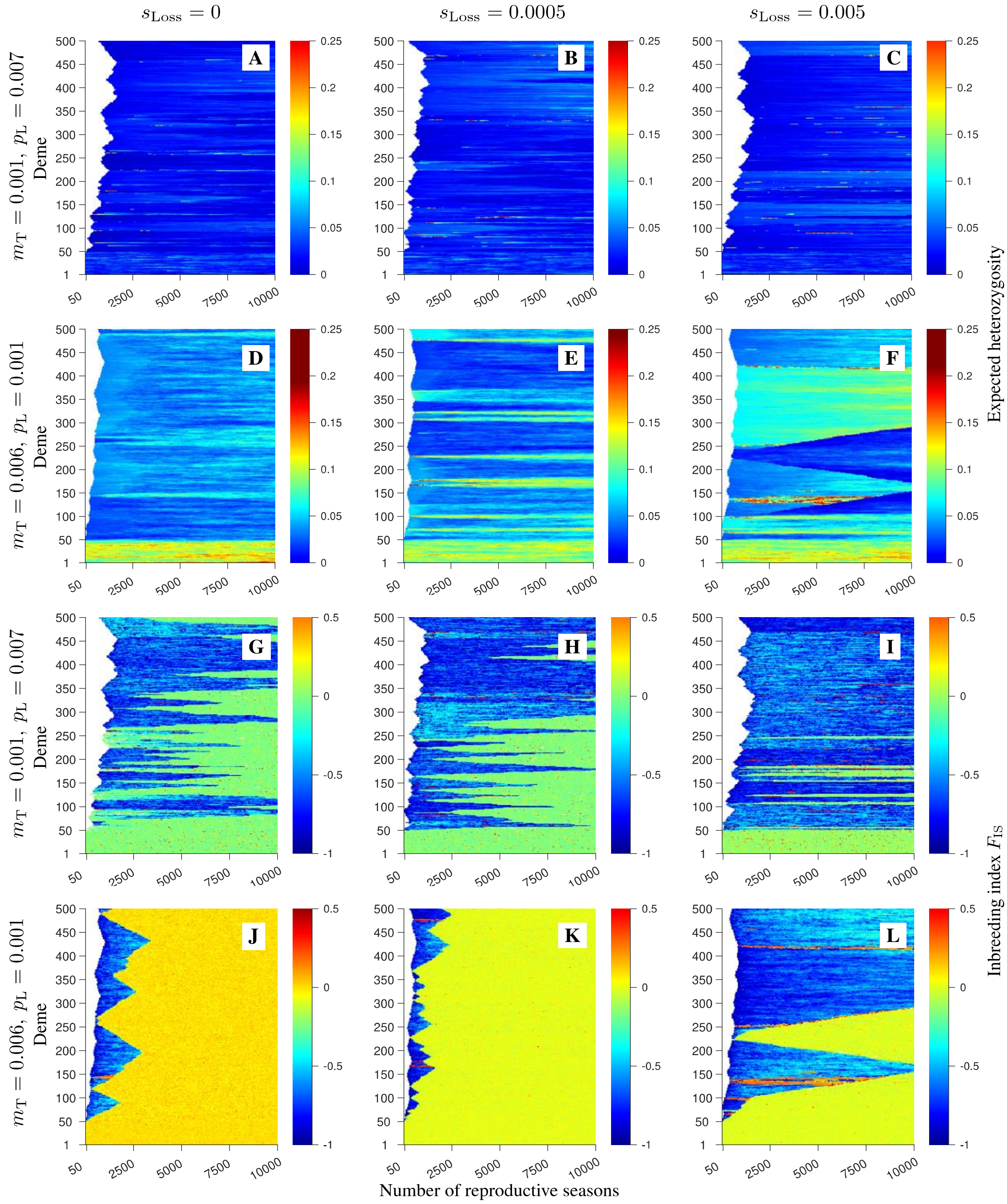

**Fig. S3.** Spatio-temporal patterns of expected heterozygosity and $\boldsymbol{F}_{\mathrm{IS}}$ obtained in single stochastic realisations of the model with both short- and long-range dispersal. The corresponding clonal structure and observed heterozygosity are shown in Fig. 4 in the main text. For the explanation of the parameters used, refer to Fig. 4 in the main text.

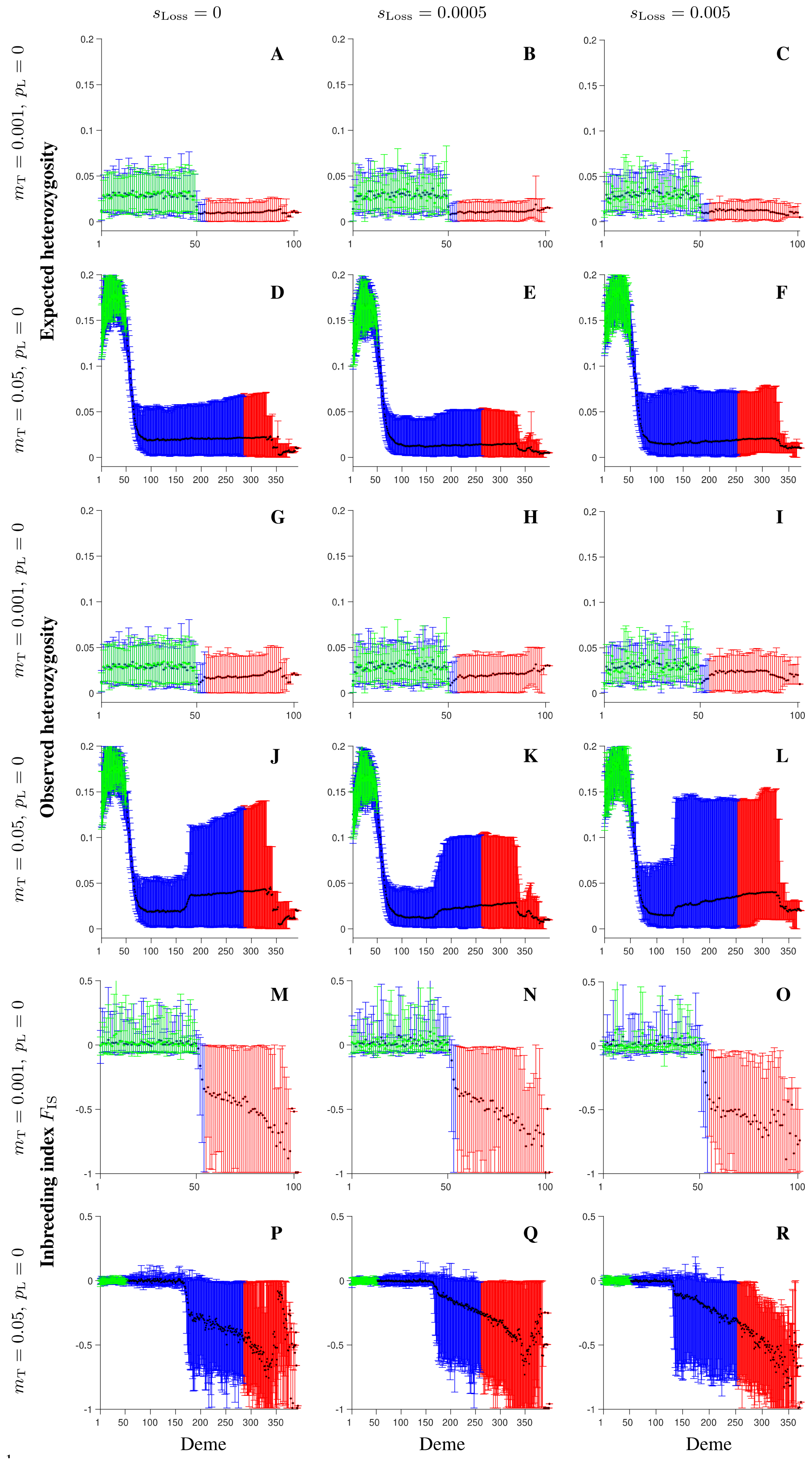

**Fig. S4.**

Expected heterozygosity (A-F), observed heterozygosity (G-L), and $F_{\mathrm{IS}}$ (M-R) obtained 1000 reproductive seasons after the start of expansion in the model where dispersal occurs only to the nearest neighbours. The black symbols show the values averaged over 100 independent realisations, the blue and red bars span between the 5th and 95th percentiles of the realised distributions in the sexual and clonal areas, respectively. For comparison, the green symbols and error bars show the corresponding results at the end of the burn-in simulations wherein only demes 1,2,…, 50 were fully occupied and the remainder of the habitat was empty. The rows (columns) differ by the value of the migration (loss of sex) rate, as depicted in the figure. Remaining parameters: total number of demes $K = 500$, maximum number of individuals in each deme $N = 100$, death rate $d = 0.1$, number of mating attempts per fully sexually active male $S = 8$, number of clonal offspring per inidivudal $C = 1$, number of loci $L = 100$, mutation rate $\mu = 0.000025$, free recombination between any pair of loci, probability of long-range dispersal $p_{\mathrm{L}} = 0$, mean long-range dispersal distance was 50 demes.. The burn-in was run for 250,000 reproductive seasons.

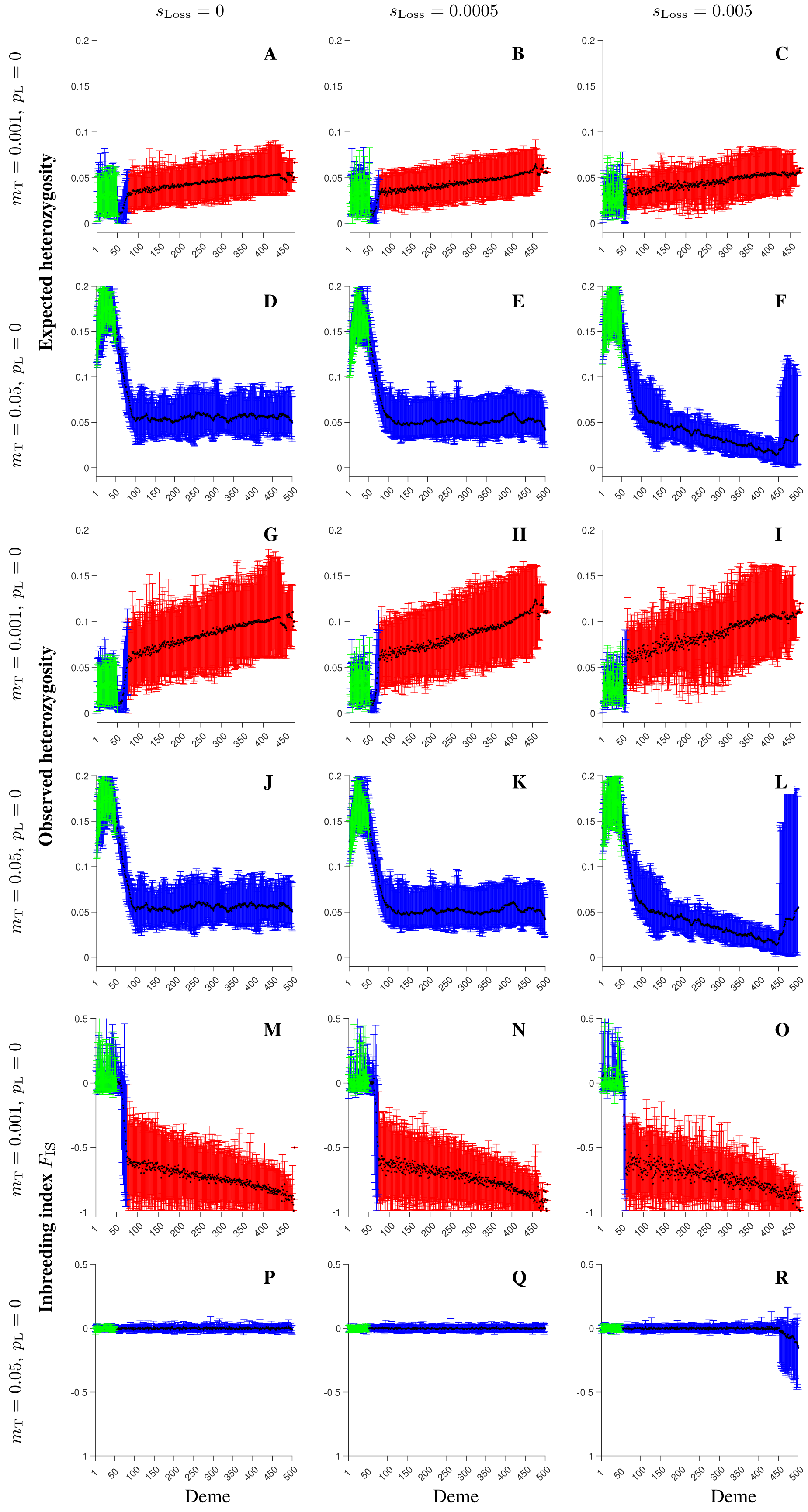

**Fig. S5.** Expected heterozygosity (A-F), observed heterozygosity (G-L), and $\boldsymbol{F}_{\mathrm{IS}}$ (M-R) obtained 10,000 reproductive seasons after the start of expansion in the model where dispersal occurs only to the nearest neighbours. The black symbols show the values averaged over 100 independent realisations, the blue and red bars span between the 5$^{th}$ and 95$^{th}$ percentiles of the realised distributions in the sexual and clonal areas, respectively. For comparison, the green symbols and error bars show the corresponding results at the end of the burn-in simulations wherein only demes 1,2,…, 50 were fully occupied and the remainder of the habitat was empty. The rows (columns) differ by the value of the migration (loss of sex) rate, as depicted in the figure. Remaining parameters are as in Figure S4.

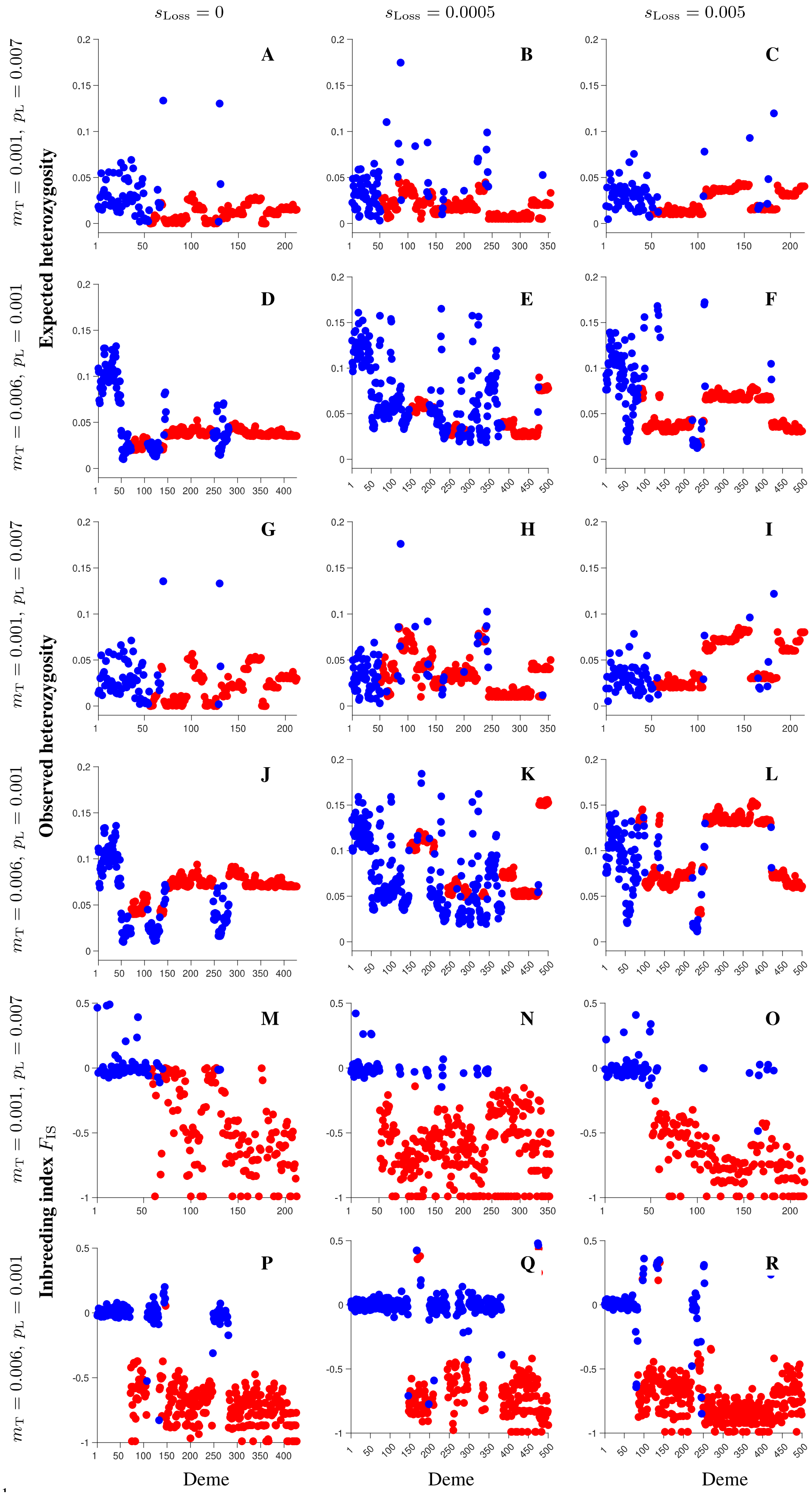

**Fig. S6**. Expected heterozygosity (top two rows), observed heterozygosity (middle two rows), and $\boldsymbol{F}_{\mathrm{IS}}$ (bottom two rows) obtained 1000 reproductive seasons after the start of expansion in the model with both short- and long-range dispersal.  The blue and red symbols depict results in sexual and clonal areas, respectively. The rows differ by the value of the total migration rate ($\boldsymbol{m}_{\mathrm{T}}$) and the probability that a migrant disperses by long-range dispersal ($\boldsymbol{p}_{\mathrm{L}}$) as depicted in the figure. The columns differ by the value of the loss of sex rate ($\boldsymbol{s}_{\mathrm{Loss}}$) as indicated in the figure. Remaining parameters are as in Figure S4.

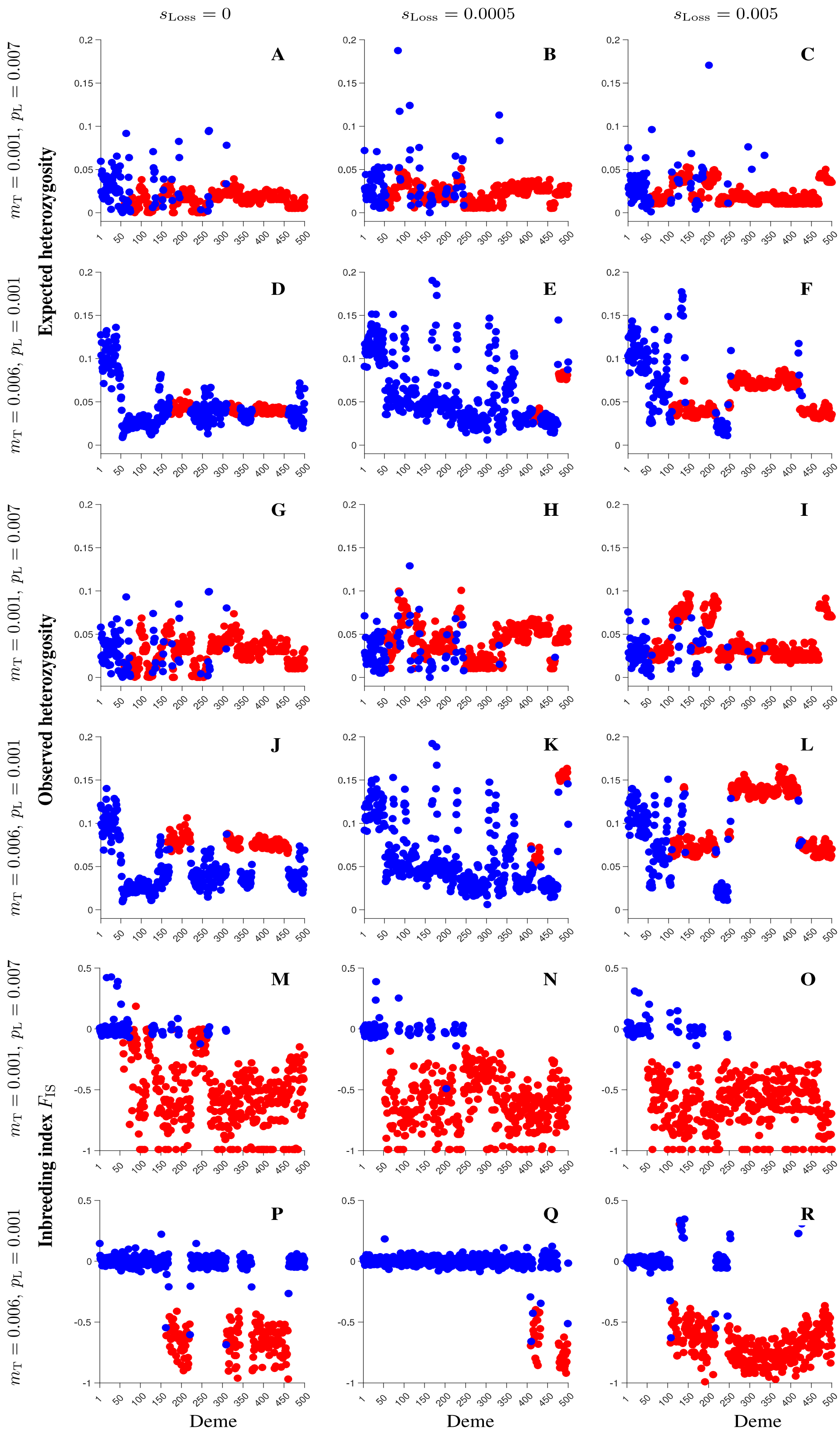

**Fig. S7**. Same as in Figure S6, but for the results obtained 2000 reproductive seasons after the start of expansion.

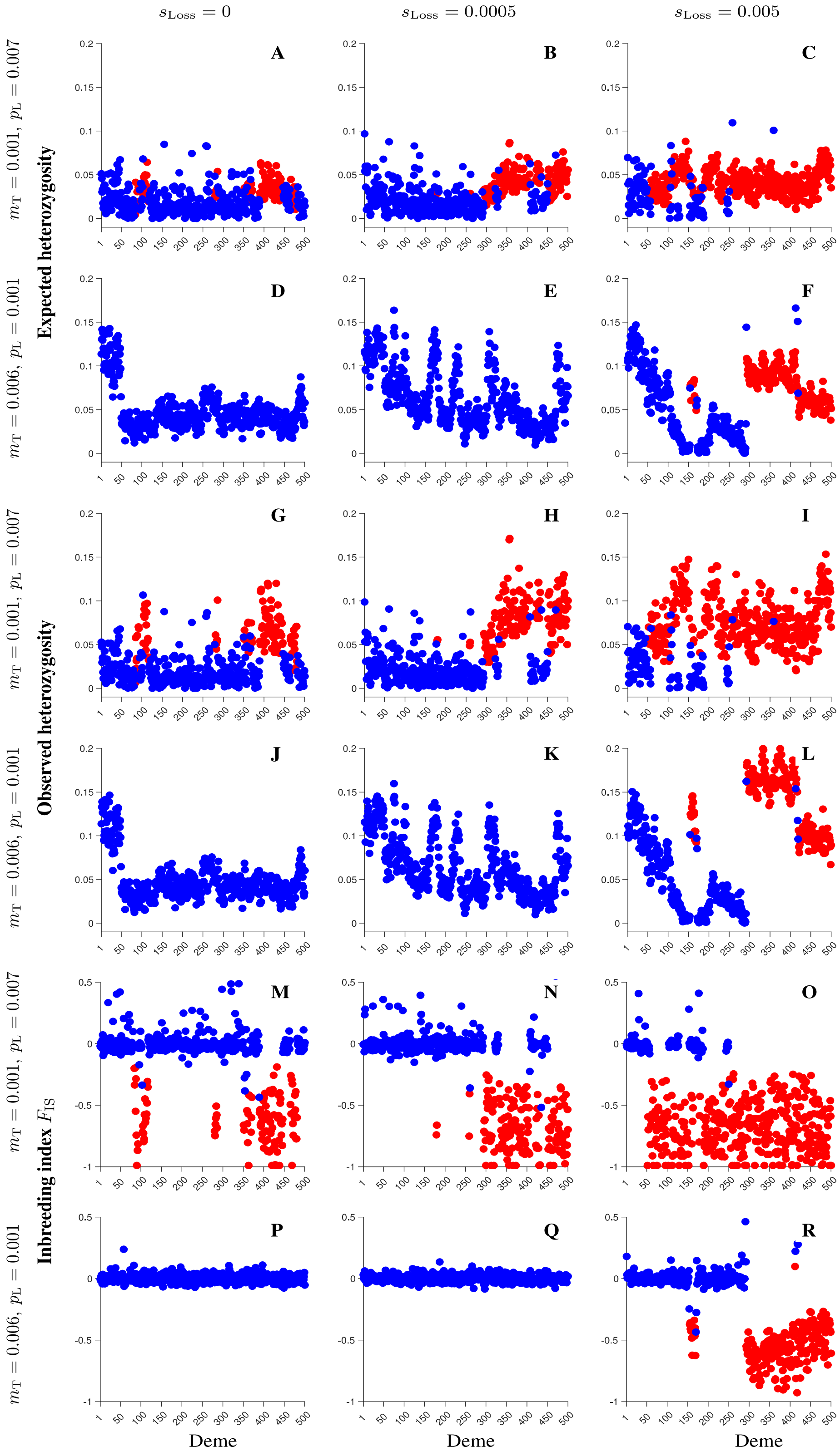

**Fig. S8**. Same as in Figures S6, but for the results obtained 10,000 reproductive seasons after the start of expansion.

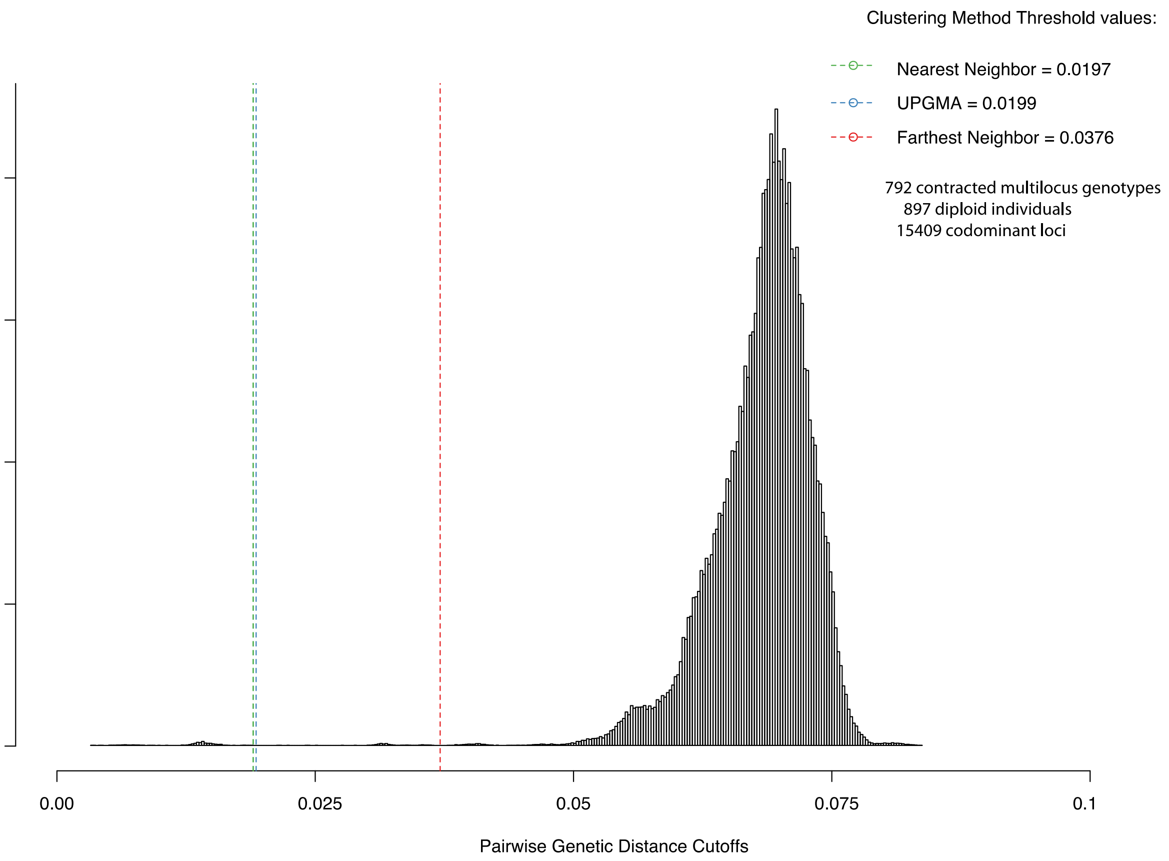

Fig. S9. Frequency distribution of genetic distances between samples using three different clustering algorithms ("Nearest Neighbour", "UPGMA" and "Farthest Neighbour"; Green, blue and red vertical dotted lines, respectively) to infer the genetic distance threshold separating asexually recruited genotypes (clones). Vertical coloured dotted lines indicate the estimated cut-off genetic distances to separate clones from sexual individuals.

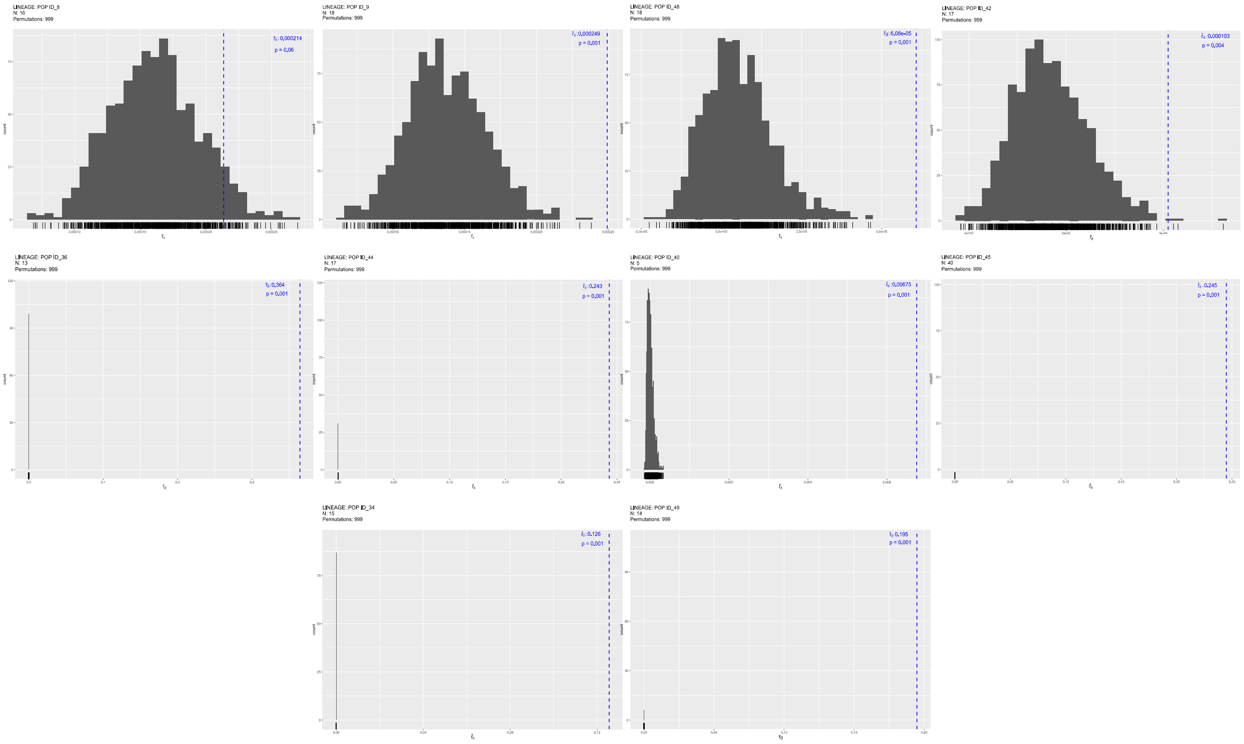

Fig. S10. Plots of the standardised index of association (*r*d) to test for linkage disequilibrium due to clonal reproduction (clones in second and third rows). Plots for each clonal lineage, with 2 additional sexual populations (Pop ID 8, 9) from the Atlantic and 2 from the Baltic (Pop ID 48, 42) (top row). *p*-value after 999 permutations show significance of deviations from linkage equilibrium for panmictic populations (See Methods).

**Table S1.** List of sites.

| Country | Area | Locality | Short name | Long | Lat | Lineage ID | N |
|---|---|---|---|---|---|---|---|
| France | Atlantic Ocean | Roscoff | | 3.990306 W | 48.72569 | 1 | 20 |
| UK | Atlantic Ocean | Bangor | | 4.130942 W | 53.23381 | 2 | 18 |
| Norway | Atlantic Ocean | Sandestrand | | 5.582986 | 59.01569 | 3 | 9 |
| Norway | Atlantic Ocean | Skadbergsanden | | 5.911303 | 58.45658 | 4 | 18 |
| Norway | Atlantic Ocean | Østhossel | | 6.668383 | 58.07486 | 5 | 10 |
| Norway | Atlantic Ocean | Lillesand | | 8.3534 | 58.2242 | 6 | 18 |
| Denmark | Atlantic Ocean | Sylt | | 8.43892 | 55.02092 | 7 | 20 |
| Sweden | Atlantic Ocean | Tjärnö | | 11.13124 | 58.86678 | 8 | 16 |
| Sweden | Atlantic Ocean | Kristineberg | | 11.44767 | 58.24773 | 9 | 14 |
| Sweden | Atlantic Ocean | Espevik | | 12.18487 | 57.18986 | 10 | 20 |
| Denmark | Transition zone | Bønnerup | | 10.71214 | 56.53311 | 11 | 20 |
| Sweden | Transition zone | Vejbystrand | | 12.76233 | 56.31566 | 12 | 17 |
| Denmark | Transition zone | Ebeltoft | | 10.66798 | 56.19467 | 13 | 17 |
| Denmark | Transition zone | Helsingør | | 12.53644 | 55.954 | 14 | 16 |
| Denmark | Transition zone | Juelsminde | | 10.00844 | 55.71683 | 15 | 18 |
| Denmark | Transition zone | Kolding | | 9.62606 | 55.51372 | 16 | 20 |
| Denmark | Transition zone | Høruphav | | 9.8855 | 54.90556 | 17 | 19 |
| Germany | Transition zone | Kiel | | 10.18825 | 54.41169 | 18 | 20 |
| Germany | Transition zone | Neustadt | | 10.80459 | 54.09126 | 19 | 20 |
| Denmark | Transition zone | Stege | | 12.17081 | 54.98386 | 20 | 19 |
| Sweden | Transition zone | Falsterbokanalen | | 12.93411 | 55.41305 | 21 | 15 |
| Sweden | Baltic Proper | Kivik | | 14.23059 | 55.68783 | 22 | 20 |
| Sweden | Baltic Proper | Ottenby (Öland) | | 16.40004 | 56.19481 | 23 | 14 |
| Sweden | Baltic Proper | Borgholm (Öland) | | 16.72077 | 56.88585 | 24 | 19 |
| Sweden | Baltic Proper | Västervik | | 16.73196 | 57.6953 | 25 | 19 |
| Latvia | Baltic Proper | Tuja | | 24.37959 | 57.48248 | 26 | 20 |
| Estonia | Baltic Proper | Kõiguste (Saaremaa) | | 22.98222 | 58.37028 | 27 | 20 |
| Estonia | Baltic Proper | Panga Pank (Saaremaa) | | 22.29861 | 58.56889 | 28 | 20 |
| Estonia | Baltic Proper | Pulli Pank (Saaremaa) | | 22.95372 | 58.61473 | 29 | 20 |
| Estonia | Baltic Proper | Sarve (Hiiumaa) | | 23.06278 | 58.84361 | 30, 31 | – |
| Sweden | Baltic Proper | Österäs | | 18.99462 | 59.70227 | 32 | 16 |
| Finland | Baltic Proper | Hankö | | 23.14083 | 59.83222 | 33 | 20 |
| Finland | Gulf of Bothnia | Skärgårdshavet | SKR | 21.83467 | 60.291 | **34**, 35 | – |
| Sweden | Gulf of Bothnia | Djursten | DJU | 18.40115 | 60.3687 | **36, 45** | – |
| Finland | Gulf of Bothnia | Rauma | | 21.30495 | 61.14348 | 37 | 20 |
| Finland | Gulf of Bothnia | Björneborg | | 21.34905 | 61.48 | 38 | 31 |
| Sweden | Gulf of Bothnia | Kuggören | KUG | 17.51643 | 61.69878 | 39, **45** | – |
| Finland | Gulf of Bothnia | Sälskäret | SAL | 21.2154 | 62.33361 | **40**, 41, **45** | – |
| Finland | Gulf of Bothnia | Storskäret | | 21.13582 | 62.47456 | 42 | 17 |
| Sweden | Gulf of Bothnia | Barstahamn | | 18.40207 | 62.861 | 43 | 17 |
| Finland | Gulf of Bothnia | South Vallgrund | VAL | 21.38194 | 63.33694 | **44, 45** | – |

| | | | | | | | |
|---|---|---|---|---|---|---|---|
| Sweden | Gulf of Bothnia | Järnäs | JAR | 19.66628 | 63.43556 | **45** | 8 |
| Finland | Gulf of Bothnia | Hällkalla | HAL | 21.08962 | 63.30737 | **45** | 7 |
| Estonia | Gulf of Finland | Pakrineeme | | 24.13362 | 59.36889 | 46 | 20 |
| Finland | Gulf of Finland | Helsinki | | 24.91765 | 60.14057 | 47 | 20 |
| Estonia | Gulf of Finland | Letipea | | 26.61055 | 59.55205 | 48 | 17 |
| Russia | Gulf of Finland | Primorsk | PRI | 28.63007 | 60.36555 | **49** | 17 |

**Table S2.** Tests of observed heterozygosity ($H_o$) and genetic variation ($H_e$) among the three geographic areas, Atlantic, Transition zone and Baltic Sea.

| Statistic | Atlantic | Transition | Baltic Sea | OSx | P-value |
|---|---|---|---|---|---|
| *All ramets included* | | | | | |
| **Het Obs** | 0.147 | 0.162 | 0.137 | 0.031 | 0.01 |
| **Het Exp** | 0.146 | 0.156 | 0.132 | 0.03 | 0.01 |
| *Ramets excluded* | | | | | |
| **Het Obs** | 0.147 | 0.162 | 0.137 | 0.031 | 0.01 |
| **Het Exp** | 0.146 | 0.156 | 0.137 | 0.024 | 0.01 |
